# Evolutionary diversification of lipid logistics shapes synaptic maturation in primates

**DOI:** 10.64898/2026.07.08.734953

**Authors:** V. Rava, E. Restelli, F. Mirabella, L. Silvestrini, E. Cannone, K. Ciuba, M. Abbas, A. Graziadei, M. Francolini, A. Pękowska, E. Taverna

## Abstract

The prolonged developmental trajectory of the human brain (neoteny) is a defining feature of human evolution. Yet, the cellular mechanisms underlying this delay remain poorly understood. Here, comparing human and chimpanzee induced neurons and cerebral organoids, we we show that human neurons form fewer excitatory synapses and synchronize network activity later. Electron microscopy further revealed reduced synaptic vesicle docking and clustering near release sites, identifying altered presynaptic assembly as a prominent feature of human neuronal development. Unexpectedly, human neurons accumulate more, not less, membrane lipids, revealing a dissociation between lipid abundance and synaptic maturation. Transcriptomics and synaptosome proteomics resolve this paradox: chimpanzee neurons preferentially engage lipid metabolism and synaptic-maturation programs, whereas human neurons upregulate intracellular lipid transport and trafficking pathways. Together, our findings identify membrane organization as a previously unrecognized regulatory layer that controls neuronal neoteny. These results also suggest that evolutionary divergence can arise through changes in the spatial deployment of membrane lipids rather than their abundance, providing a molecular framework for understanding the evolution of human brain neoteny.

## INTRODUCTION

The prolonged developmental trajectory of the human brain is a defining feature of our species evolution and is thought to contribute to the extended plasticity and computational capacity of human neural circuits. Comparative studies of humans and great apes have revealed species-specific differences in developmental timing across multiple stages of corticogenesis, from neural progenitors to differentiated neurons and neural circuits (1–8). Among these, delayed neuronal maturation, referred to as neoteny, has emerged as a recurring hallmark of the human lineage, yet the cellular mechanisms underlying this heterochrony remain poorly understood (4,5).

Stem cell-derived neuronal systems provide an experimentally tractable platform to compare neurodevelopmental processes across primate species (1,9,10). In particular, induced neurons generated through forced expression of Neurogenin 2 (NGN2) reproduce key aspects of neuronal maturation and have revealed intrinsic differences between human and chimpanzee neurons, including delayed acquisition of synaptic and network activity in humans (11–13). These findings suggest that species-specific differences in maturation are encoded within neuronal programs themselves, but the molecular and cell biological mechanisms responsible remain largely unresolved (12).

Synapse formation is a major determinant of neuronal maturation. The establishment of functional circuits requires coordinated assembly of presynaptic release sites, postsynaptic specializations and activity-dependent signaling mechanisms (14–17). Even modest alterations in the timing or efficiency of these processes can profoundly affect circuit development. Human-specific changes in synaptic regulators, including SRGAP2 paralogs, have implicated synaptogenesis as a key substrate for evolutionary modification of neuronal connectivity (18–20). However, it remains unclear whether delayed maturation reflects differences in synapse abundance, ultrastructure, molecular composition or membrane organization.

Membrane lipids are increasingly recognized as central regulators of synaptic function. Cholesterol- and glycosphingolipid-enriched membrane domains influence vesicle docking, neurotransmitter release, receptor organization and synaptic plasticity (17,21–32). These processes are tightly regulated through neuron–glia interactions, which coordinate lipid synthesis, trafficking and redistribution across cellular compartments (23–27). Recent work has highlighted the importance of lipid metabolism and membrane organization for synaptic physiology (22,25,27), yet whether species-specific differences in lipid handling contribute to the distinct maturation trajectories of human and chimpanzee neurons has not been explored.

Here, we use a combination of comparative primate stem cell models, brain organoids, and multi-scale structural, molecular and functional analyses to investigate the cellular and molecular basis of human neuronal neoteny. We show that delayed synaptic maturation in human neurons is accompanied by coordinated alterations in synaptic ultrastructure, membrane organization and lipid-associated molecular programs. Together, these findings suggest that evolutionary differences in neuronal maturation are associated with the spatial organization of membrane lipids rather than with changes in lipid abundance alone.

## RESULTS

### Human and chimpanzee induced neurons provide a comparative model of neuronal maturation

To investigate cellular mechanisms underlying species-specific differences in neuronal maturation, we generated induced neurons (iNs) from previously established human and chimpanzee induced pluripotent stem cell (iPSC) lines through doxycycline-inducible expression of NGN2 (Fig. 1A,B). Neuronal differentiation was monitored for up to 35 days using molecular, structural and functional analyses.

**Figure 1.**
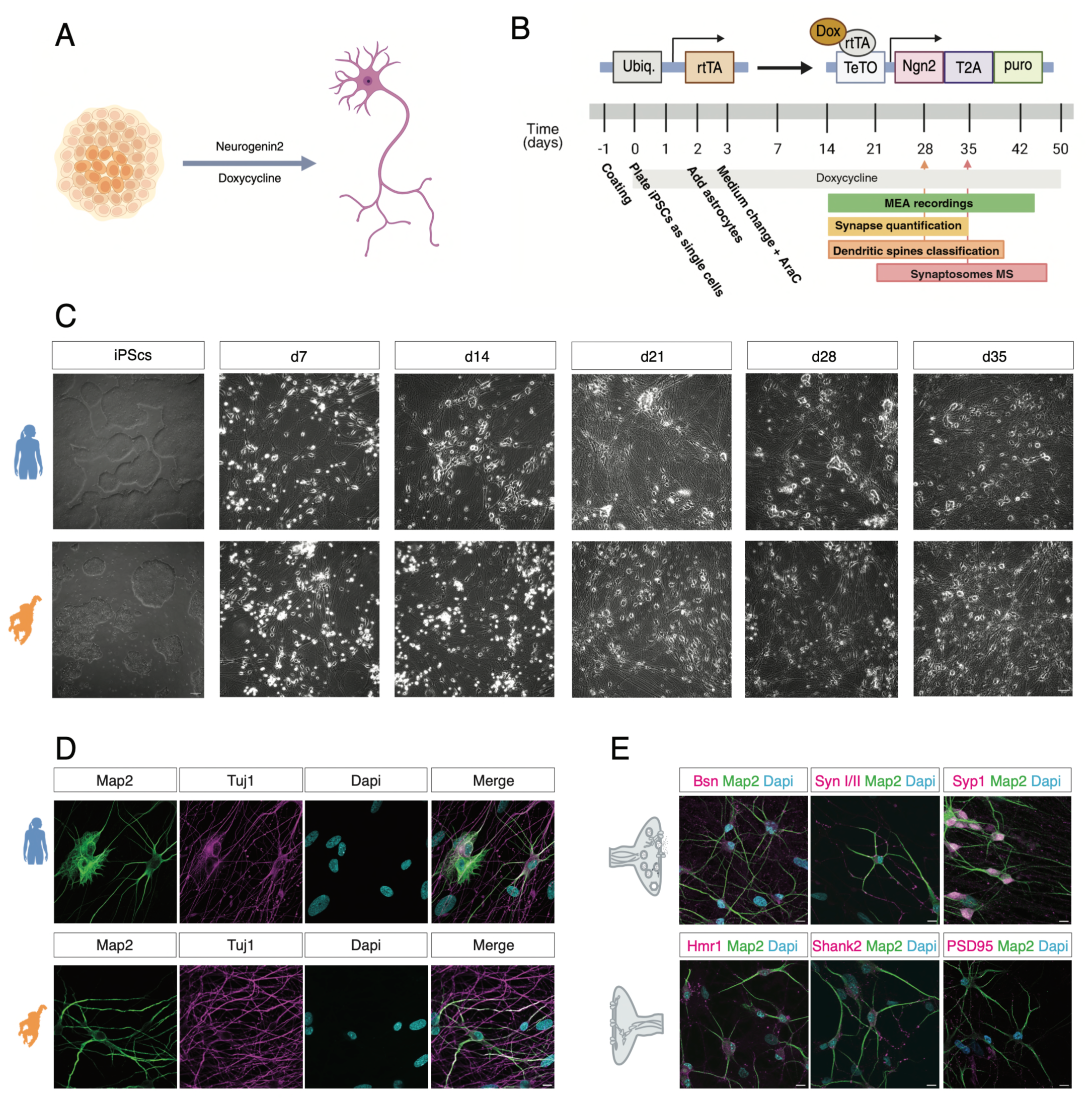
Human and ape induced neurons enable comparative neuronal cell biology across evolution. (A) Schematic overview of neuronal differentiation. Induced pluripotent stem cells (iPSCs) were differentiated into induced neurons (iNs) through Neurogenin2 (Ngn2) overexpression upon doxycycline induction. (B) Schematic representation of Ngn2 vector (top), experimental timeline of neuronal differentiation and downstream analyses (bottom). Following Ngn2 induction, neuronal cultures were analyzed over time (d7–d50), with assays including MEA recordings, synapse quantification, dendritic spine analysis, synaptosome proteomics, and electron microscopy. (C) Brightfield images showing morphological progression of human (top) and chimpanzee (bottom) cultures during differentiation from iPSCs to iNs across time points (iPSCs, d7, d14, d21, d28, d35). Scale bars: iPSCs inset 200µm, iNs inset 50µm. (D) Immunofluorescence characterization of neuronal identity. Human iNs express neuronal cytoskeletal markers as MAP2 and Tuj1, confirming neuronal differentiation, scale bars are 10µm. (E) Immunofluorescence staining of synaptic markers expression. Pre-synaptic markers (Bassoon, Synapsin I/II, Synaptophysin) and post-synaptic markers (Homer1, Shank2, PSD-95) are detected along MAP2-positive neurites, indicating the formation of synaptic structures, scale bars are 10µm.

Both human and chimpanzee iNs extended elaborate neurite networks within the first week of differentiation and formed dense neuronal cultures by day 14 (Fig. 1C). Immunofluorescence analysis confirmed expression of neuronal markers including MAP2 and TUJ1, as well as progressive accumulation of pre- and postsynaptic proteins including Bassoon, Synapsin, Synaptophysin, Homer1, Shank2 and PSD95 (Fig. 1D,E).

### Human neurons show delayed maturation of network activity

To assess functional maturation, we performed longitudinal recordings using multi-electrode arrays (MEAs) between days 21 and 42 of differentiation (Fig. 2A). NeuroFluor labeling revealed progressive elaboration of neuritic networks in both species (Fig. 2B). Despite comparable network formation, functional activity differed markedly. Chimpanzee cultures developed synchronized bursting activity during maturation, whereas human neurons displayed predominantly asynchronous activity throughout the recording period (Fig. 2C). Consistent with these observations, chimpanzee cultures exhibited a greater number of active channels at later stages of differentiation (Fig. 2D).

**Figure 2.**
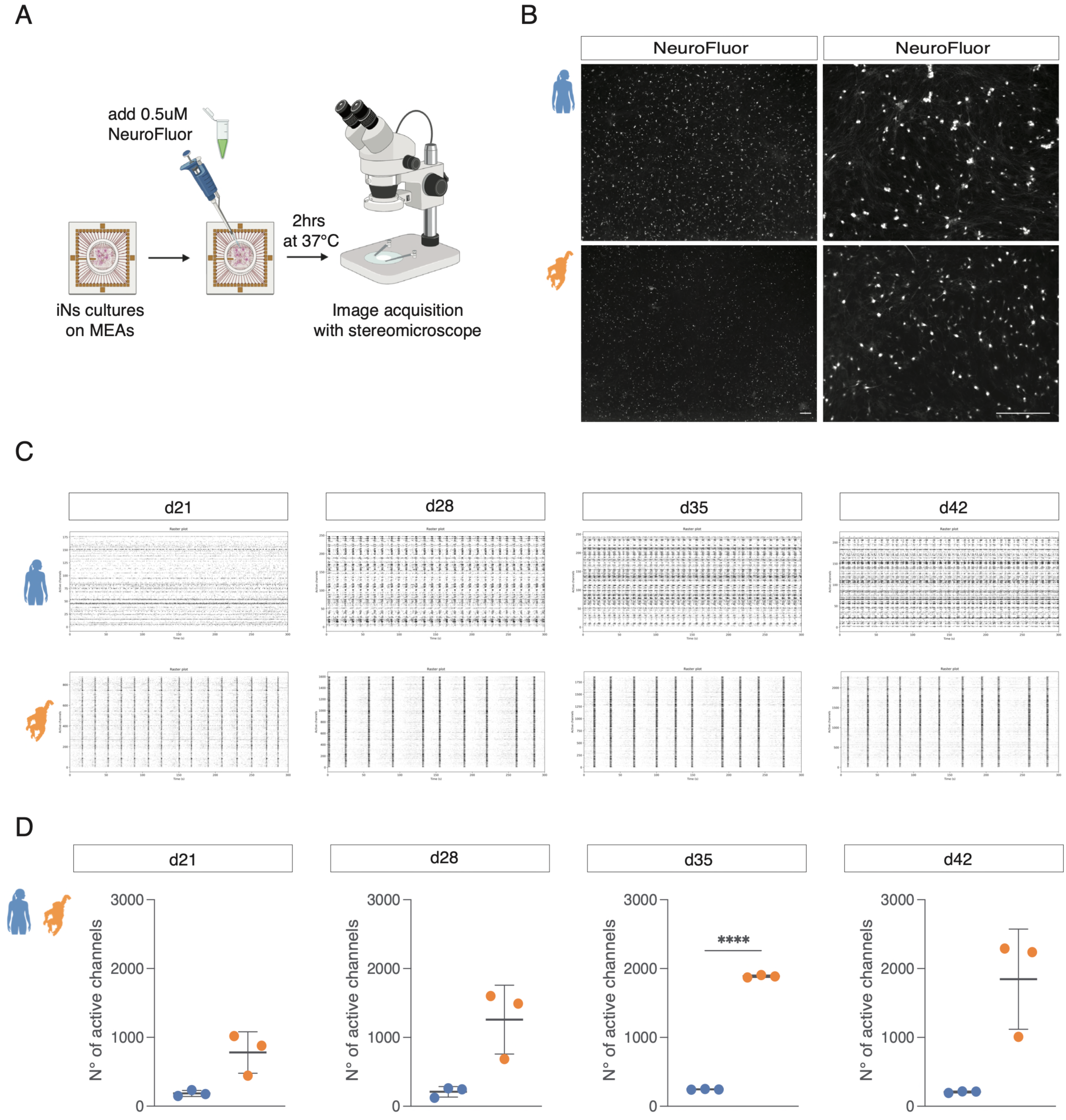
Delayed maturation of network activity in human neurons. (A) Schematic representation of the NeuroFluor workflow used to assess cell density and cell distribution on the MEA chip. (B) Epifluorescence images of human and chimpanzee neurons at d14, after incubation with NeuroFluor, scale bars are 100µm, and 50µm. (C) Raster plots that represent spontaneous network activity in human and chimpanzee iNs over time (d21, d28, d35, d42) of only active channels. n= number of MEA wells/batches: Human n=3/1, Chimpanzee n=3/1. (D) Quantification, using welch t test, of the number of active channels in each chip comparing the two species over time (d21, d28, d35, d42). n= number of MEA wells/batches: Human n=3/1, Chimpanzee n=3/1. Each dot represents a MEA chip; statistical significance was determined using Welch’s t test, bars indicate mean ± [SD]. ****P < 0.0001.

### Human neurons form fewer synaptic contacts

To determine whether delayed network maturation was associated with altered synaptogenesis, we quantified pre- and postsynaptic puncta along sparsely labeled dendrites using Bassoon and Homer1 immunostaining (Fig. 3A–C). Both species formed detectable synaptic puncta throughout maturation; however, chimpanzee neurons exhibited significantly higher densities of pre- and postsynaptic structures and increased numbers of colocalized puncta at later stages (Fig. 3D,E). Similar results were obtained using an independent combination of synaptic markers (Supplementary Fig. 3). These data indicate reduced synapse formation in human neurons.

**Figure 3.**
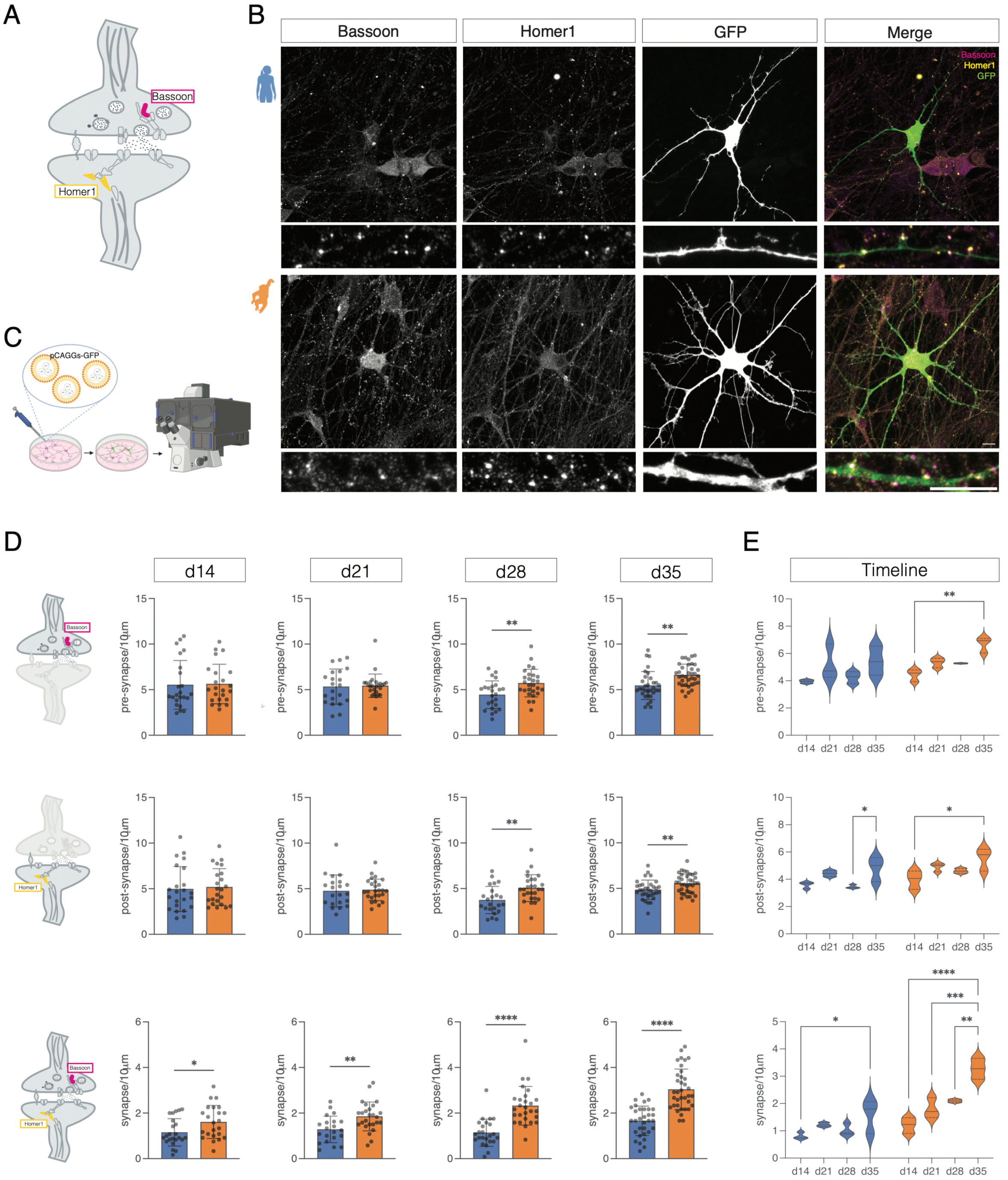
Reduced synapse formation in human neurons during functional maturation. (A) Schematic representation of excitatory synapses showing the localization of the presynaptic marker Bassoon and the postsynaptic marker Homer1 used for synapse quantification. (B) Representative immunofluorescence images of human (top) and chimpanzee (bottom) induced neurons (iNs) stained for Bassoon (presynaptic), Homer1 (postsynaptic), and GFP to visualize sparsely labeled neurons. Merged images show the distribution and colocalization of pre- and postsynaptic puncta along dendrites. Insets highlight dendritic segments used for quantification. Scale bars 10µm. (C) Experimental scheme illustrating sparse single-cell labeling using pCAGGs-GFP plasmid to enable visualization of individual neuronal morphology and accurate quantification of synaptic puncta along isolated dendritic arbors. (D) Quantification of synaptic puncta density across developmental time points (d14, d21, d28, d35). Density of presynaptic (Bassoon⁺), postsynaptic (Homer1⁺), and colocalized synaptic puncta (synapses) is shown per 10µm of dendrite. Chimpanzee neurons exhibit a progressive increase in synaptic density over time, whereas human neurons show reduced pre- and postsynaptic puncta density and significantly fewer colocalized synaptic puncta at later stages (d28–d35). Each dot represents a neuron. A total of 5-10 neurons were analyzed for each replicate, 4 technical replicates were performed, 1 biological replicate. Statistical significance was determined using Welch’s t test, bars indicate mean ± [SD]. (E) Summary of synaptic maturation across time. Violin plots show the distribution of presynaptic, postsynaptic, and colocalized synaptic puncta densities per 10µm of dendrite across developmental stages. Chimpanzee neurons display increased synapse density and maturation over time, whereas human neurons exhibit a delayed trajectory of synapse formation. A total of 5-10 neurons were analyzed for each replicate, 4 technical replicates were performed, 1 biological replicate. Statistical significance was determined using two-way ANOVA (multiple comparisons). *P < 0.05, **P < 0.005, ***P < 0.0005, ****P < 0.0001.

### Dendritic spine architecture is largely conserved between species

We next examined dendritic spine organization using sparse GFP labeling and super-resolution imaging (Fig. 4A). In contrast to synapse density, overall dendritic spine density did not differ significantly between species (Fig. 4B). Classification of dendritic protrusions into filopodia, long thin, thin, stubby and mushroom categories revealed similar distributions in human and chimpanzee neurons (Fig. 4G). Both species exhibited predominantly immature filopodia-like protrusions.

**Figure 4.**
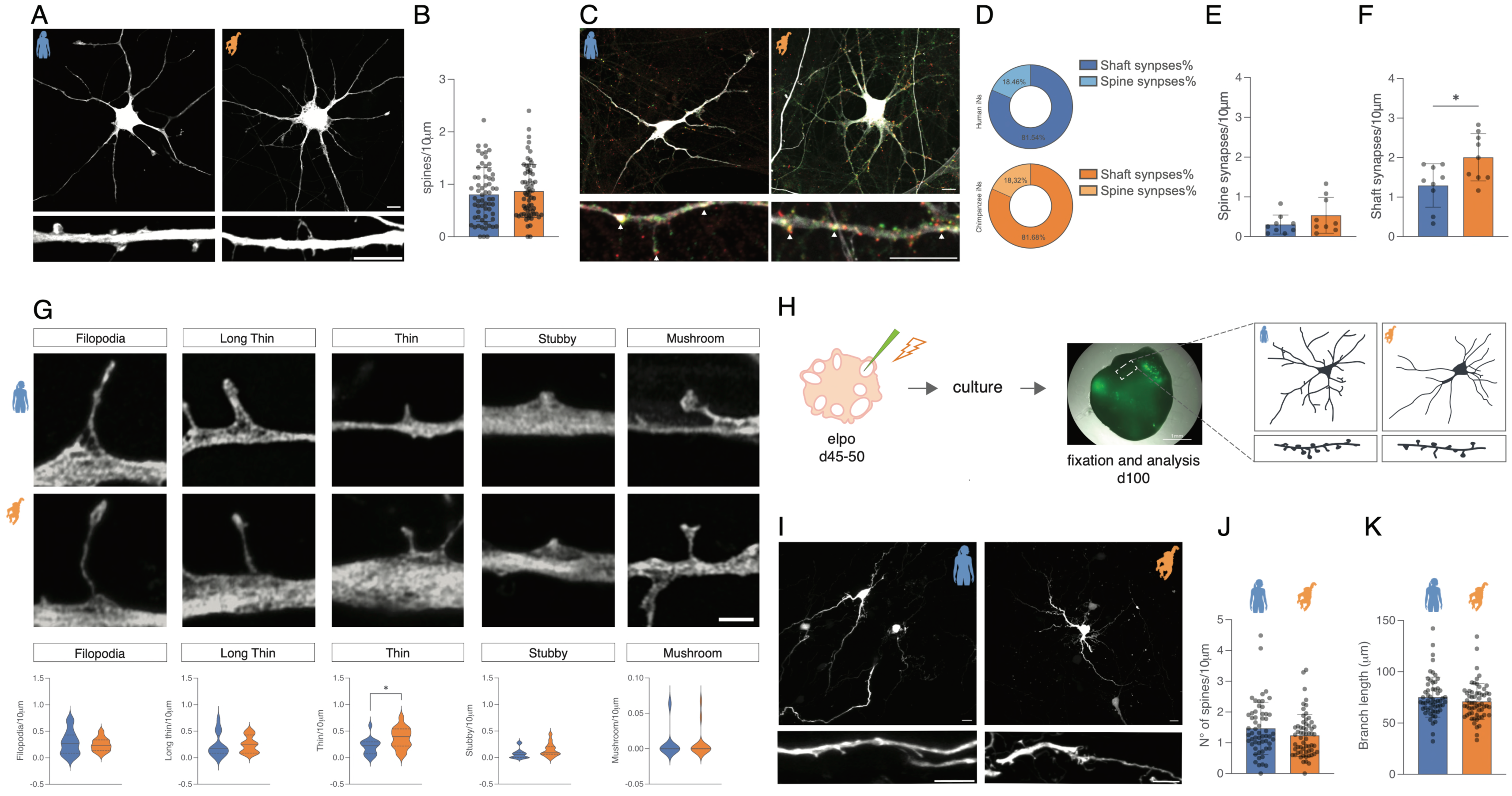
Architecture of human and chimpanzee dendritic spines in induced neurons and brain organoids. (A) Representative images of sparsely GFP labeled human (blue) and chimpanzee (orange) induced neurons (iNs) showing dendritic morphology and high-magnification views of dendritic segments used for spine analysis. Scale bars 10µm. (B) Quantification of dendritic spine density per 10µm of dendrite in human and chimpanzee neurons. Human and chimpanzee neurons exhibit the same spine density compared at matched developmental stages. A total of 5-11 neurons were analyzed for each replicate, 3 technical replicates were performed, 1 biological replicate. Statistical significance was determined using Welch’s t test, bars indicate mean ± [SD]. (C) Representative images of synapse localization along dendrites, distinguishing spine synapses and shaft synapses in human and chimpanzee neurons. Scale bars 10µm. (D) Pie chart representing the percentage of spine synapses and shaft synapses in human (blue) and chimpanzee (orange). (E) Quantification of the number of spine synapses every 10µm in human and chimpanzee iNs A total of 3 neurons were analyzed for each replicate, 3 technical replicates were performed, 1 biological replicate. Statistical significance was determined using Welch’s t test, bars indicate mean ± [SD]. (F) Quantification of the number of shaft synapses every 10µm in human and chimpanzee iNs A total of 3 neurons were analyzed for each replicate, 3 technical replicates were performed, 1 biological replicate. Statistical significance was determined using Welch’s t test, bars indicate mean ± [SD]. *P < 0.05. (G) Classification of dendritic spines morphology. Representative images of spine types, including filopodia, long thin, thin, stubby, and mushroom spines (top), and quantification of their relative frequency per 10µm of dendrite length (bottom). Human and chimpanzee neurons display an equal proportion of immature and mature spine types. Scale bar, 2µm. A total of 5-11 neurons were analyzed for each replicate, 3 technical replicates were performed, 1 biological replicate. Statistical significance was determined using Welch’s t test, bars indicate mean ± [SD]. *P < 0.05. (H) Experimental workflow for dendritic spine analysis in human brain organoids. Organoids were electroporated (elpo) at day 45–50, followed by culture, fixation, and analysis to assess neuronal morphology and spine development at d100. (I) Representative images dendritic spines in human and chimpanzee organoid derived neurons. Scale bars 10µm. (J) Quantification of dendritic spines density in human and chimpanzee organoid-derived neurons. Human and chimpanzee neurons display the same spine density. A total of 18-20 neurons were analyzed for each replicate, 3 technical replicates were performed, 1 biological replicate. Statistical significance was determined using Welch’s t test, bars indicate mean ± [SD]. (K) Quantification of branch length (in 10µm), indicator of dendritic complexity, in human and chimpanzee organoids derived neurons. A total of 18-20 neurons were analyzed for each replicate, 3 technical replicates were performed, 1 biological replicate. Statistical significance was determined using Welch’s t test, bars indicate mean ± [SD].

To extend these observations and mimic the three-dimensional context of developing brain, we analyzed GFP-labeled neurons in human and chimpanzee cerebral organoids (Fig. 4H–K). Consistent with two-dimensional cultures, spine density was comparable between species. Most synapses were localized to dendritic shafts, whereas only a minority were associated with dendritic spines. The relative distribution of shaft and spine synapses was similar between species despite the reduced overall number of synaptic contacts in human neurons (Fig. 4C–F).

### Human synapses show reduced presynaptic vesicle coupling

To examine synaptic organization at ultrastructural resolution, we performed transmission electron microscopy on differentiating neuronal cultures (Fig. 5A,B). Synaptic boutons were present in both species; however, chimpanzee neurons displayed greater clustering of synaptic vesicles and a larger proportion of docked vesicles at presynaptic terminals (Fig. 5C,D,G,H). Human synapses contained fewer vesicles positioned near the active zone and exhibited reduced vesicle docking, indicating weaker spatial coupling between vesicle pools and release sites.

**Figure 5.**
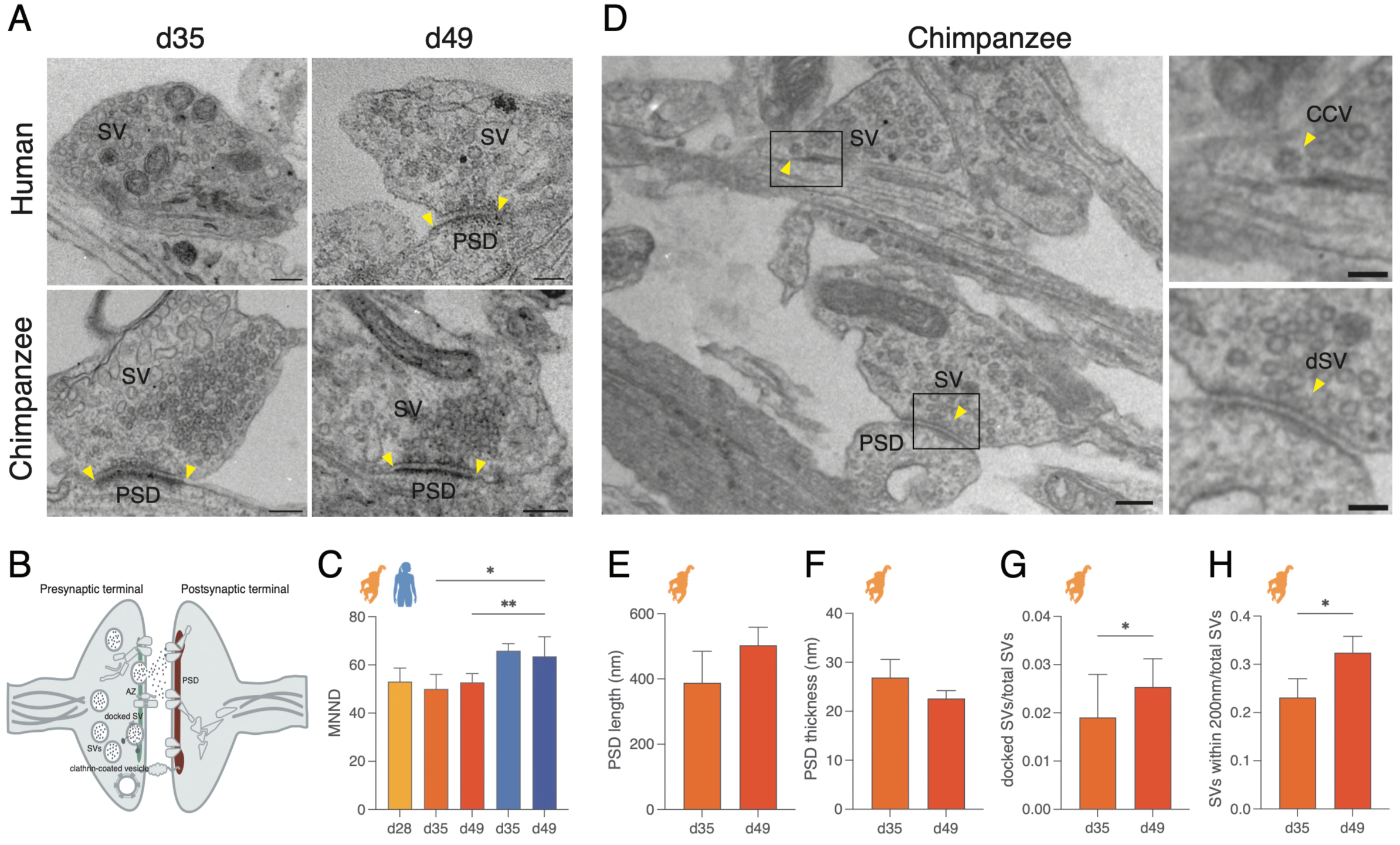
Synapse ultrastructure suggests lower coupling of synaptic vesicles with the release sites in human neurons. (A) Ultrastructural analysis of synapses. Representative transmission electron microscopy images of human and chimpanzee iNs at d35 and d49 showing synaptic vesicles (SVs), postsynaptic densities (PSD). Scale bars 200nm. (B) Schematic representation of a chemical synapse, highlighting the quantified ultrastructural features: synaptic vesicles (SVs), clathrin-coated vesicles, docked SV, active zone (AZ), postsynaptic density (PSD). (C) Quantification of MNND in chimpanzee iNeurons d28, d35, and d49 and in human iNeurons d35 and d49. A total of 4-41 synapses were analyzed for each replicate, 1-5 technical replicates were performed, 1 biological replicate. Statistical significance was determined using two-way ANOVA (multiple comparisons). *P < 0.05, **P < 0.005. (D) Images showing ultrastructure hallmarks of functional synapses in chimpanzee iNeurons including clathrin-coated vesicles (CCV, inset 1) and docked synaptic vesicles (dSV, inset 2). CCV and dSV are indicated by arrowheads. Scale bars: 200nm (left), 100nm (insets). (E, F) Quantification of PSD length and PSD thickness in chimpanzee iNeurons at d35 and d49. A total of 11-14 synapses were analyzed for each replicate, 1-5 technical replicates were performed, 1 biological replicate. Statistical significance was determined using Mann-whitney test (PSD length, and Unpaired t test (PSD thickness). (G) Quantification of docked synaptic vesicles in chimpanzee iNeurons at d35 and d49, normalized to the total of the synaptic vesicles per synapse. A total of 15-23 synapses were analyzed for each replicate, 1-5 technical replicates were performed, 1 biological replicate. Statistical significance was determined using Mann-whitney test. *P < 0.05. (H) Quantification of synaptic vesicles within 200nm of the active zone in chimpanzee iNeurons at d35 and d49, normalized to the total synaptic vesicles per synapse. A total of 15-23 synapses were analyzed for each replicate, 1-5 technical replicates were performed, Statistical significance was determined using Mann-whitney test. *P < 0.05.

Postsynaptic density (PSD) parameters were quantified primarily in chimpanzee neurons because relatively few mature PSDs were detected in human cultures. Although PSD length and thickness did not change significantly during maturation, PSD thickness showed a tendency to increase over time (Fig. 5E,F).

These observations highlight altered presynaptic organization as a prominent ultrastructural feature of human neurons, indicating impaired coupling between synaptic vesicles and the release machinery.

### Human neurons exhibit altered GM1-associated membrane-domain organization

Efficient synaptic vesicle docking and neurotransmitter release require the coordinated organization of the presynaptic membrane. Cholesterol- and glycosphingolipid-enriched membrane domains regulate vesicle trafficking, neurotransmitter release and the assembly of presynaptic protein complexes. One such membrane component is, the ganglioside GM1, which is widely used as a marker of ordered membrane domains and can be detected with high specificity using Cholera Toxin Subunit B (CTXB). During neuronal development, lipids are supplied largely by astrocytes, allowing the support of neuronal membrane biogenesis and synapse formation. We therefore examined whether species-specific differences in GM1-associated membrane domains accompany the delayed presynaptic maturation observed in human neurons (Fig. 6).

**Figure 6.**
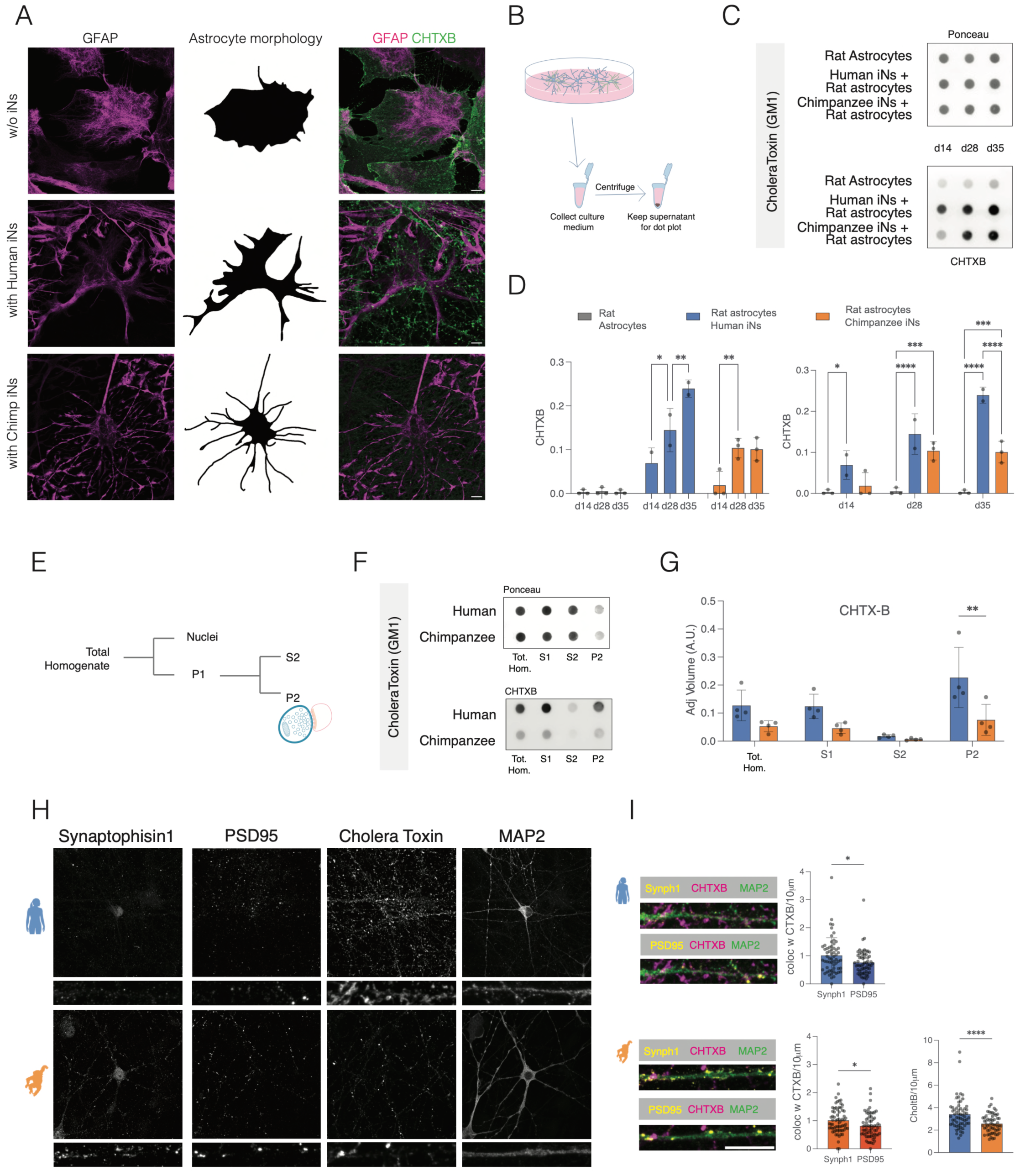
Human neurons exhibit altered GM1-associated membrane domain organization and neuron–astrocyte lipid interactions. (A) Representative images (left and right panel) of rat astrocytes cultured alone (top) or co-cultured with human (middle) or chimpanzee (bottom) induced neurons (iNs), stained for the astrocytic marker GFAP, and Choleratoxin subunit B (CHTXB). Astrocytes undergo a transition from a flat morphology in monoculture to a branched morphology in the presence of neurons. Scale bars 10µm. Schematic representation (middle panel) of astrocyte morphological changes observed in mono-culture and neuron–astrocyte co-culture conditions. (B) Experimental workflow for collection and analysis of conditioned medium from astrocyte and neuron–astrocyte co-cultures. (C) Representative dot blot analysis of conditioned medium using cholera toxin B subunit (CTXB) to detect GM1-associated membrane domains. Ponceau staining is shown as loading control and used for CHTXB normalization. (D) Quantification of CHTXB signal in conditioned medium collected from rat astrocytes cultured alone or co-cultured with human or chimpanzee iNs at different developmental stages (d14, d28 and d35). 2-3 technical replicates were performed, 1 biological replicate. Statistical significance was determined using two-way ANOVA (multiple comparisons). *P<0.05, **P<0.005, ***P<0.0005, ****P<0.0001. (E) Schematic representation of biochemical fractionation used to isolate synaptosome-enriched fractions (P2) from neuronal cultures. (F) Representative dot blot analysis of total homogenate (Tot.), post-nuclear supernatant (S1), cytosolic fraction (S2), and synaptosome-enriched fraction (P2) from human and chimpanzee neuronal cultures. Ponceau staining is shown as loading control. (G) Quantification of CHTXB signal across cellular fractions. Human neuronal cultures show increased GM1-associated signal in synaptosome-enriched fractions compared to chimpanzee cultures. 4 technical replicates were performed, 1 biological replicate. Statistical significance was determined using two-way ANOVA (multiple comparisons). **P<0.005. (H) Representative immunofluorescence images of human (top) and chimpanzee (bottom) neurons stained for Synaptophysin1, PSD95, CHTXB, and MAP2. Higher magnification views of dendritic segments are shown below. Scale bars 10µm. (J) Quantification of CHTXB colocalization with presynaptic (Synaptophysin1) and postsynaptic (PSD95) markers, and quantification of CHTXB signal normalized to dendritic length. CHTXB-positive domains show partial overlap with synaptic markers and are increased in human neurons relative to chimpanzee neurons. Scale bars 10µm. A total of 50-60 dendrites were analyzed for each replicate, 1 technical replicate were performed, 1 biological replicate. Statistical significance was determined using Welch’s t test, bars indicate mean ± [SD]. *P<0.05, ****P<0.0001.

Rat astrocytes cultured alone exhibited the expected flattened morphology, whereas co-culture with neurons induced a branched, stellate morphology in both human and chimpanzee cultures, indicating comparable neuron–astrocyte interactions under both conditions (Fig. 6A,B). We next quantified GM1 using CTXB. Notably, conditioned medium from human neuronal cultures contained significantly more GM1 than medium from chimpanzee cultures (Fig. 6C,D). Consistent with this finding, biochemical fractionation showed increased GM1-associated signal in synaptosome-enriched fractions prepared from human neurons relative to chimpanzee neurons (Fig. 6E–G).

To determine how increased amount of GM1 was distributed within neurons, we examined CTXB-labelled membrane domains relative to pre- and postsynaptic markers. CTXB-positive membrane domains were detected throughout neuronal processes and showed only partial overlap with both Synaptophysin and PSD95. Thus, despite their enrichment in synaptosome-containing fractions, GM1-associated membrane domains were not selectively confined to mature synaptic sites but were broadly distributed along neurites in both species (Fig. 6H,I).

Altogether, these findings reveal an unexpected dissociation between GM1 abundance and synaptic maturation. Although human neurons accumulated higher levels of GM1 in both conditioned medium and synaptosome-enriched fractions, they exhibited reduced synapse formation and impaired presynaptic vesicle docking. Likewise, the distribution of GM1-associated membrane domains relative to synaptic markers was similar across species, suggesting that increased GM1 abundance is not accompanied by preferential accumulation at synaptic sites.

These data suggest that membrane lipid abundance alone is insufficient to support presynaptic maturation and raise the possibility that species differences instead arise from the differences in membrane remodelling and presynaptic assembly between species. To explore this possibility, we next profiled the protein composition of synaptosome-enriched fractions from human and chimpanzee neurons.

### Synaptosome proteomics identifies species-specific membrane and lipid-associated pathways

To identify molecular pathways associated underlying interspecies differences, we performed quantitative proteomic analysis of synaptosome-enriched fractions isolated from human and chimpanzee neurons (Fig. 7A). Fractionation efficiency was confirmed by enrichment of Synaptophysin, PSD95 and GluA1 in the synaptosomal fraction (Fig. 7B–D).

**Figure 7.**
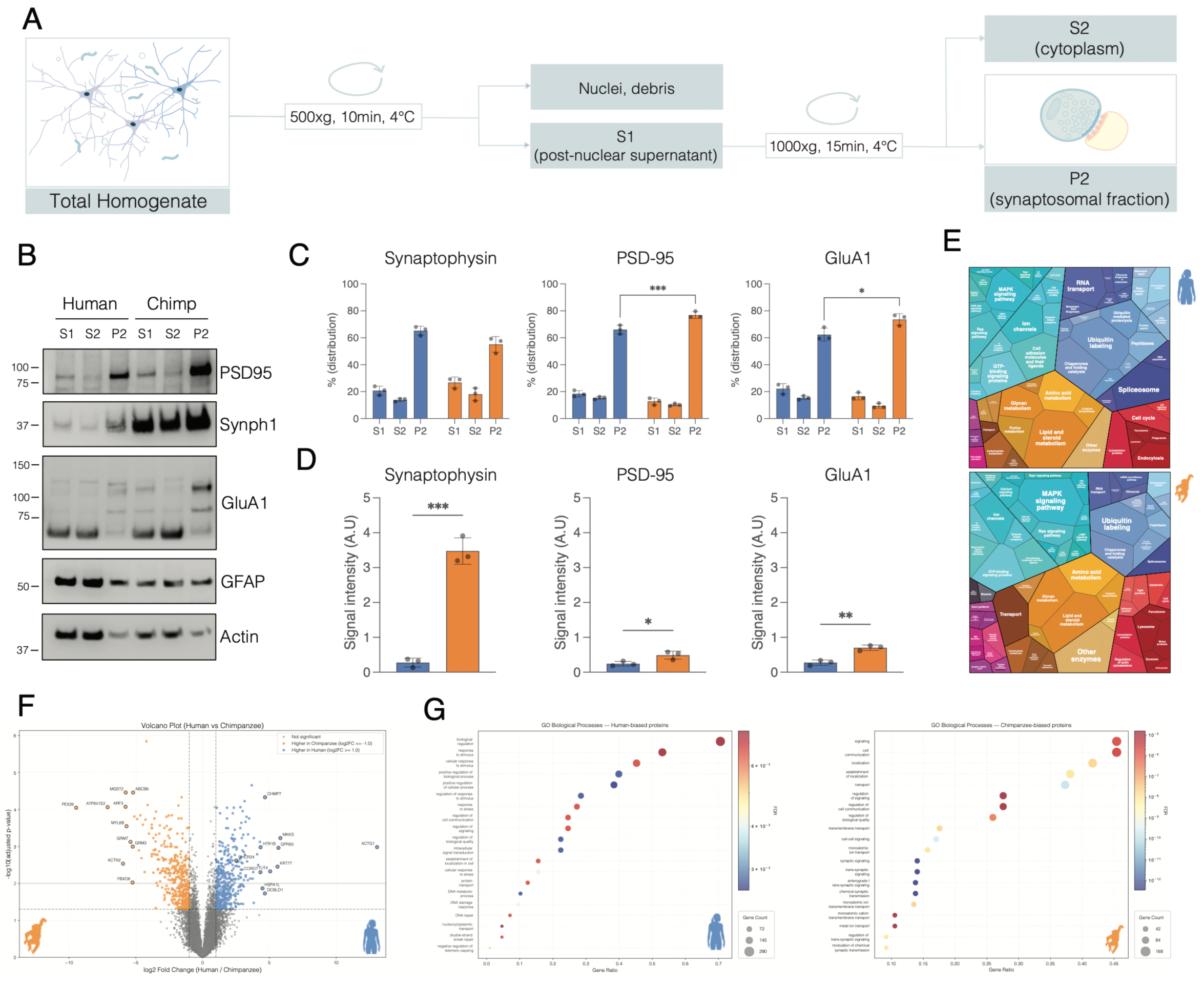
Comparative subcellular proteomics reveals species-specific differences in synaptic membrane and lipid-associated pathways. (A) Schematic overview of synaptosome preparation. Neuronal cultures were homogenized and subjected to differential centrifugation to obtain post-nuclear supernatant (S1), cytosolic fraction (S2), and synaptosome-enriched fraction (P2). (B) Immunoblot analysis of fractionation efficiency in human and chimpanzee samples. Synaptic proteins (PSD-95, Synaptophysin, GluA1) are enriched in the synaptosomal (P2) fraction, whereas astrocytic (GFAP) and cytoskeletal (Actin) markers show distinct distribution across fractions. (C) Quantification of protein enrichment across fractions (S1, S2, P2) for Synaptophysin, PSD-95, and GluA1, confirming enrichment of synaptic proteins in the P2 fraction. A total of 3 technical replicates were performed, 1 biological replicate. Statistical significance was determined using two-way ANOVA (multiple comparisons), bars indicate mean ± [SD]. *P<0.05, ***P<0.0005. (D) Quantification of synaptic protein levels in the P2 fraction. Chimpanzee samples show increased levels of synaptic proteins (Synaptophysin, PSD-95, GluA1) compared to human samples. A total of 3 technical replicates were performed, 1 biological replicate. Statistical significance was determined using Welch’s t test, bars indicate mean ± [SD]. *P<0.05, **P<0.005, ***P<0.0005. (E) Proteomap representation of the synaptosomal proteome in human (up) and chimpanzee (down) neurons, illustrating the relative contribution of functional categories, including membrane organization, intracellular transport, signaling pathways, and metabolic processes. Mass spectrometry proteomics was performed on 5 individual technical replicates, 1 biological replicate. (F) Volcano plot of differential protein abundance between human and chimpanzee synaptosomes. Each point represents a protein plotted by fold change (1) and statistical significance (alpha=0.05). Proteins enriched in human (orange) and chimpanzee (blue) samples highlight species-specific differences in synaptic molecular composition. Mass spectrometry proteomics was performed on 5 individual technical replicates, 1 biological replicate. (G) Functional enrichment analysis of differentially abundant proteins. Gene Ontology (GO) and pathway analyses show that chimpanzee-enriched proteins are associated with synaptic signaling, trans-synaptic communication, and ion transport, whereas human-enriched proteins are associated with intracellular transport, cellular stress responses, and regulation of biological processes.

Proteomic profiling identified 8,690 proteins across samples and revealed extensive species-specific differences in synaptic composition (Fig. 7E,F). Proteins enriched in chimpanzee synaptosomes were associated with synaptic signaling, trans-synaptic communication and ion transport, whereas human synaptosomes were enriched for pathways related to intracellular transport, cellular regulation and stress responses (Fig. 7G).

Several differentially abundant proteins converged on membrane organization and lipid handling, including ARF1, CHMP7, CORO7, ATP6V0D2, ATAD3B, APOL2 and DHCR24. These proteins are associated with vesicle trafficking, endolysosomal processing, membrane remodeling and sterol metabolism, suggesting coordinated differences in membrane logistics between species.

We next asked whether distinct protein composition, underlying the differences in neuronal development could be associated with neurodevelopmental diseases. Indeed, disease-enrichment analysis revealed that proteins predominantly found in human synaptosomes significantly overlap with gene networks associated with neurological and psychiatric disorders. In contrast, chimpanzee synaptosome-enriched proteins were predominantly associated with synaptic signaling and neuronal function (Supplementary Fig. 4). These observations indicate that molecular pathways distinguishing human and chimpanzee synapses also intersect with networks implicated in human brain disease.

### Human and chimpanzee neurons engage distinct lipid-handling programs

Our proteomic analysis revealed extensive species-specific differences in the molecular composition of developing synaptosomes, particularly among proteins involved in membrane trafficking, vesicle dynamics and synaptic organization. Yet, whether these molecular differences reflected local properties of developing synapses or broader neuronal maturation programs remained unclear. To address this question, we compared the transcriptomes of human and chimpanzee neurons across neuronal maturation to determine whether species-specific differences in synaptic composition are accompanied by coordinated changes in global gene expression (Fig. 8A-D, Methods).

**Figure 8.**
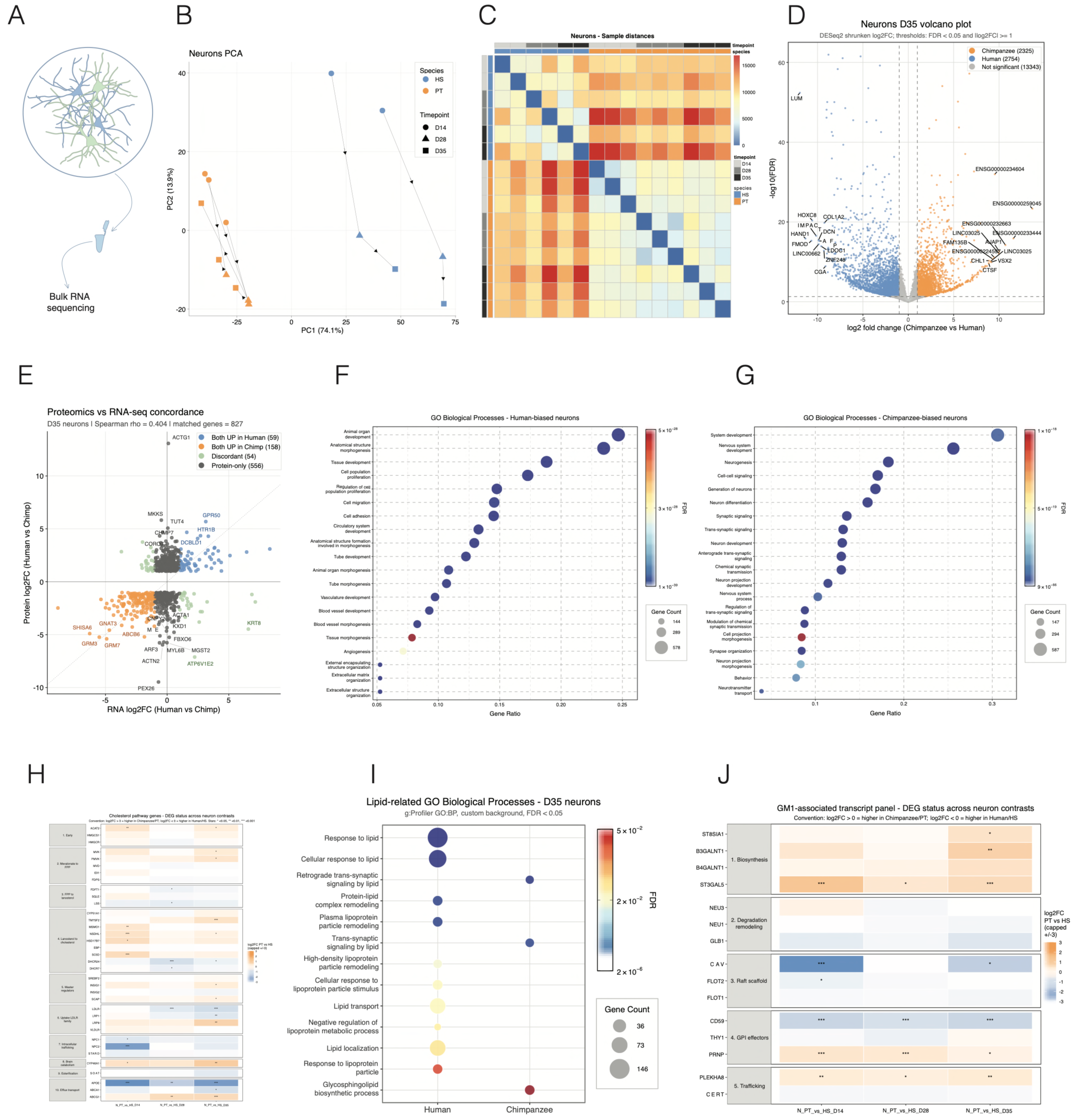
Human and chimpanzee neurons engage distinct transcriptional lipid programs. (A) Experimental overview of bulk RNA sequencing analysis performed on human and chimpanzee iNs collected at different stages of maturation. (B) Principal component analysis (PCA) of transcriptomic profiles from human and chimpanzee neurons at d14, d28, and d35. Samples segregate by both species and developmental stage, revealing distinct maturation trajectories. Bulk RNA sequencing was performed on 2 (human) and 3 (chimpanzee and rat astrocytes mono-cultures) individual technical replicates, 1 biological replicate. (C) Sample-to-sample distance heatmap showing clustering of transcriptomic profiles across developmental stages and species. Replicates cluster together and separate according to species identity and maturation state. (D) Volcano plot of differentially expressed genes between human and chimpanzee neurons at d35. Genes enriched in human neurons (blue) and chimpanzee neurons (orange) are shown according to fold change and statistical significance. (E) Comparison of differential expression detected by bulk RNA sequencing (d35) and synaptosome proteomics at d35. Scatter plot shows concordance between transcript and protein abundance changes. Genes identified as significantly enriched at both RNA and protein levels in human or chimpanzee neurons are highlighted. (F) Gene Ontology (GO) biological process enrichment analysis of genes preferentially expressed in human neurons. Enriched terms include cellular transport, migration, extracellular organization, and developmental regulatory pathways. (G) GO biological process enrichment analysis of genes preferentially expressed in chimpanzee neurons. Enriched categories include neuronal maturation, synaptic signaling, neurotransmitter transport, and nervous system development. (H) Heatmap showing differential expression of genes associated with cholesterol-related pathways, including cholesterol transport, trafficking, and regulatory processes. Expression values are shown across developmental stages and species. (I) Enrichment analysis of lipid-related GO biological processes. Human neurons show preferential enrichment of pathways associated with lipoprotein handling, plasma lipoprotein particle organization, lipid transport, and protein–lipid complex remodeling. (J) Heatmap showing differential expression of genes associated with GM1- and glycosphingolipid-related pathways. While biosynthetic pathways show limited species-specific differences, genes involved in trafficking, transport, and pathway regulation are preferentially enriched in human neurons. Together, these analyses reveal distinct transcriptional programs underlying lipid handling in human and chimpanzee neurons, with chimpanzee neurons preferentially enriched for neuronal maturation and lipid metabolic pathways, and human neurons enriched for lipid transport, trafficking, and membrane regulatory processes. Data are presented as mean ± SEM where applicable. Differential expression analysis was performed using DESeq2 with Benjamini–Hochberg correction for multiple testing. GO enrichment analyses were performed using g:Profiler. Bubble size indicates gene count and color indicates adjusted significance values (FDR).

Principal component analysis separated samples according to both developmental stage and species, indicating distinct transcriptional trajectories during neuronal maturation (Fig. 8B,C). Differential expression analysis identified 2,461 genes enriched in human neurons and 2,060 genes enriched in chimpanzee neurons at day 35 of differentiation (Fig. 8D, Methods). Comparison of transcriptomic and synaptosome proteomic datasets revealed overlapping species-biased molecular signatures, confirming that transcriptional differences are reflected in the molecular composition of developing synapses (Fig. 8E).

Functional enrichment analysis of genes differentially expressed between human and chimpanzee iNeurons at day 35 revealed distinct maturation programs being activated in the two species. Consistent with our previous observations, chimpanzee neurons preferentially expressed genes associated with neuronal maturation and synaptic signalling, indicating a more advanced neuronal maturation program. In contrast, human neurons were enriched for pathways related to cell adhesion, intracellular transport, lipoprotein handling and lipid transport (Fig. 8F–I). A striking difference emerged within lipid-associated pathways: chimpanzee neurons preferentially expressed genes involved in lipid metabolism, whereas human neurons upregulated genes regulating lipoprotein handling and intracellular lipid transport. Importantly, this divergence was not driven by differences in lipid biosynthesis. Expression of enzymes involved in cholesterol and glycosphingolipid synthesis was broadly similar between species, whereas genes regulating intracellular lipid transport, membrane trafficking and pathway regulation exhibited substantially greater divergence (Fig. 8H,J).

Together, our findings identify a species-specific program of neuronal maturation that distinguishes human and chimpanzee neurons at multiple levels of organization. Human neurons form fewer mature synapses and exhibit impaired presynaptic vesicle docking despite increased GM1-associated membrane domains. These phenotypic differences are accompanied by coordinated changes in the molecular composition of developing synapses and by broader transcriptional programs regulating intracellular membrane trafficking, lipid transport and neuronal maturation. Collectively, these observations indicate that delayed maturation of human neurons is associated not with reduced membrane abundance, but with distinct cellular programs governing membrane remodelling and utilization during synapse development.

## DISCUSSION

Neuronal neoteny is widely considered as a defining feature of human brain evolution, yet the cellular mechanisms that regulate the prolonged maturation of human neurons remain largely unknown (4,5,33,34). By combining comparative analyses of human and chimpanzee induced neurons with ultrastructural, proteomic and transcriptomic approaches, we identify membrane lipid organization as a candidate mechanism associated with species-specific differences in synaptic maturation. Rather than implicating changes in lipid abundance alone, our data suggest that evolutionary divergence is associated with differences in the spatial deployment of membrane lipids across neuronal compartments.

Across multiple experimental modalities, human neurons consistently exhibited delayed synaptic maturation. Human cultures developed coordinated network activity more slowly, formed fewer excitatory synapses and displayed altered presynaptic ultrastructure. The most prominent phenotype was observed at the presynaptic terminal, where human synapses contained fewer docked vesicles and reduced vesicle clustering near release sites. Because vesicle docking and release critically depend on the molecular organization of the presynaptic membrane (21,29–31), these observations suggested that altered membrane organization could contribute to species-specific maturation dynamics.

Our analysis of lipid-associated membrane domains supports this possibility. Human neuronal cultures exhibited increased GM1-associated signal together with altered neuron–astrocyte interactions, despite reduced synapse density and delayed network maturation. At first glance, these findings appear inconsistent with the established role of cholesterol-rich membrane domains in promoting synapse formation and neurotransmission (21,23–27,29–31). However, they instead point to a distinction between lipid abundance and lipid organization. We propose that the critical variable is not the total amount of membrane lipids available to the neuron, but their regulated deployment into specialized membrane domains that support synaptic assembly and function.

Independent molecular analyses converged on this interpretation. Synaptosome proteomics identified coordinated differences in proteins involved in vesicle trafficking, membrane remodeling and sterol homeostasis, including ARF1, CHMP7, CORO7, ATP6V0D2, ATAD3B, APOL2 and DHCR24. Bulk transcriptomics further demonstrated that the strongest species-specific divergence occurs not in cholesterol or glycosphingolipid biosynthetic pathways, but in pathways controlling lipid trafficking, intracellular transport and membrane organization. Conversely, chimpanzee neurons preferentially engaged lipid metabolic and synaptic maturation programs. Together, these observations suggest that human and chimpanzee neurons deploy distinct lipid-handling strategies during maturation.

These findings extend current models of neuronal neoteny. Previous work has established that delayed maturation is a cell-intrinsic property of human neurons and has implicated developmental timing and synaptic regulators in this process (4,5,12,18–20). Moreover, previous studies, performed on mice models, highlight the importance of astrocyte to neuron lipid transport and especially of neurons ability to uptake and use lipids (35–37). Our data suggest an additional layer of regulation, in which membrane lipids act not only as structural components or metabolic substrates, but as spatial organizers of synaptic assembly. In this framework, neuron–astrocyte interactions regulate not simply lipid supply, but the subcellular distribution of membrane lipids, thereby influencing where and when functional synapses mature. Importantly, our findings converge with those of the companion study (Ciuba et al., *Biorxiv* 2026 Primate Astrocyte Evolution Controls the Tempo of Neuronal Development) which approaches the problem from the perspective of astrocyte evolution. Whereas our work identifies neuron-intrinsic membrane remodelling and lipid transport programs associated with delayed synaptic maturation, Ciuba et al. demonstrate that astrocytes actively regulate the tempo of neuronal development through an evolutionarily remodelled TEAD–APOE signalling axis. Together, our two studies challenge the prevailing neuron-centric view of brain neoteny by identifying complementary neuron-intrinsic and astrocyte-dependent mechanisms that converge on lipid-associated pathways. This convergence suggests that coordinated lipid regulation across the neuron–astrocyte interface represents a fundamental mechanism governing the prolonged developmental trajectory of the human brain. Our study has several limitations. Although multiple independent datasets converge on lipid-associated pathways, direct perturbation of lipid trafficking will be required to establish causality. In addition, CHTXB reports GM1-enriched membrane domains rather than cholesterol directly, and future lipidomic analyses of synaptic membranes and membrane microdomains will be required to define the lipid species responsible for the observed phenotypes. Finally, although induced neurons provide a powerful comparative system, they cannot fully recapitulate the complexity of in vivo neuron–glia interactions during cortical development.

More broadly, our findings suggest that evolutionary innovation may arise not only through changes in gene regulation or metabolic pathways, but also through changes in the spatial organization of conserved cellular processes. Although demonstrated here for membrane lipids during synaptic maturation, regulated deployment of cellular components within specialized subcellular domains may represent a general mechanism through which conserved molecular machinery generates species-specific developmental programs.

## ACKNOWLEDGEMENTS

We are grateful to the services and facilities of HT for the outstanding support provided, notably, N. Maghelli, F. Casagrande, A. Fasciani and the team of the National Facility (NF) of Light Imaging, G. Faga’ and A. Corti and the team of Genome Editing and Disease Modeling, and the team Structural Biology especially Structural Proteomics unit and Cryo-electron Microscopy unit.

We thank Carlo Besta Neurological Institute (VISION@Besta), Milan-Italy, for the access to the transmission electron microscope.

We thank M. Beltrame, M. Polenghi, G. Visani and P. Cucchi for the experimental support and all members of the Taverna lab for helpful discussions.

## AUTHOR CONTRIBUTIONS

V.R. and E.T. conceptualized the work and co-wrote the manuscript. V.R. performed all the experimental work on iNs and on organoids. F.M. and V.R. performed and analysed MEA experiments. A.G. and V.R. performed mass spec experiments. L.S. and A.G. analysed the proteomics data. L.S. and M.A. analysed the bulkSeq data with V.R.. E.R. and V.R. performed western blotting and dot blot experiments. E.C. designed, performed and quantified EM experiments under the supervision of M.F. A.P. and K.C. provided additional experimental support and discussion. V.R. is a Ph.D. student within the European School of Molecular Medicine (SEMM).

## MATERIALS AND METHODS

### iPSCs culture

Human iPSCs (409B2) were obtained from RIKEN, chimpanzee iPSCs (SandraA) were obtained from MPI-EVA, Leipzig. The rtTA/Ngn2-positive iPSCs were previously generated, following Frega et al., 2017 protocol, using lentiviral vectors to stably integrate transgenes into iPSCs genomes, as described by Schörning et al., 2021.

### Thawing iPSCs

iPSCs were thawed by putting the cryogenic vial was the waterbath at 37°C for a maximum of 2 minutes. Cells were transferred, with a wet pipet tip, into a falcon already containing 5ml of DMEM/F12 (Thermofisher, 31330038). The cell suspension was centrifuged for 5 minutes at 200rcf. After centrifugation, the supernatant was discarded, and the cell pellet was resuspended into mTeSR™1 (StemCell, 85850) with 50U/ml Pen Strep (P/S) (Thermofisher, 15140122), 50µg/ml G418 (Sigma-Aldrich, G8168), 0.5 µg/ml Puromycin (Sigma-Aldrich, P9620), and 10µM Y-27632 dihydrochloride (Rock Inhibitor, RI) (MedChemExpress, 129830-38-2). Then, 1/3 and 2/3 of cells were plated into new wells (previously coated with 0.25 ug/cm^2^ Recombinant Laminin iMatrix – 511 (AMSBIO, AMS. 892 012) with a final volume of 2ml iPSCs culture medium with RI.

### iPSCs maintenance

To induce differentiation of iPSCs into iNeurons two iPSCs lines were used 409B2_Ngn2 and SandraA_Ngn2. iPSCs were cultured on 6 well plates coated with iMatrix–511 and kept in iPSCs culture medium the culture medium was changed daily; the cells were kept in incubation at 37°C and 5% CO2. To maintain the iPSCs culture cells were passaged once a week, when somewhere about 80% of confluency was reached. To passage cells, one wash with Dulbecco’s phosphate-buffered saline (DPBS) (Thermofisher, 14190-094) was done before incubation with 1ml 0.5mM UltraPure 0.5 M EDTA (pH 8.0) (Thermofisher, 15575-038) for 5 minutes at 37°C to detach cells. EDTA was discarded and 1ml of fresh iPSCs culture medium with RI was added and used to detach cells through four to six washings to maintain intact colonies. Then the cell suspension was transferred to a new well; previously coated RI was kept in the culture medium for 24 hours. For brain organoids generation, two iPSCs lines were used one chimpanzee (SandraA), one human (WTCII). For iPSCs maintanace cells were cultured in mTeSR1, and when splitted the cells were detached with ReLeSR Enzyme-Free (StemCell 5872-# 100-0483).

### Freezing iPSCs

To freeze iPSCs, the cells were washed with DPBS and incubated for 5 minutes at 37°C with 1ml of Tryple Express Enzyme (1X) (TrypLE) (Thermofisher, 12605-010). Cells were then detached from the plate with 3 ml of DMEM/F12. The suspension was transferred to a falcon and centrifuged for 5 minutes at 200rcf. The supernatant was discarded, and the cell pellet was resuspended with 500µl of mFreSR (StemCell Technologies, 5854), transferred to a cryogenic vial and progressively frozen to -80°C in a freezing box. After 24 hours the cryovials were conserved into liquid nitrogen for long-term storage.

### iNeurons differentiation from iPSCs

iPSCs differentiation into induced neurons protocol was adapted from Frega et al., 2017 and Schörnig et al., 2021.

#### Day-1

##### Coating

Either plates or glass coverslips were beforehand sterilized with 70% ethanol and then dried under UV light for 30 minutes. Afterwards they were coated with of 50µg/ml Poly-L-Ornithine solution (PLO) (Sigma-Aldrich, P4957). Plates were incubated overnight at 37°C and 5% CO2.

#### Day 0

##### Coating and plating

PLO was discarded, and the plates were washed three times with ultrapure double distilled water (ddH2O), then 10µg/ml Laminin (Sigma-Aldrich, L2020) was added to the plate and incubated at 37°C for 2 hours. Plates were then washed with ddH2O and dried under the hood. In the meantime, to detach iPSCs from the 6 well plate, the medium was discarded and cells were washed once with DPBS. Cells were incubated with 1ml of Accutase (Sigma-Aldrich, A6964) for 5 minutes at 37°C, washed with 2ml of DMEM/F12, and transferred to a falcon. iPSCs were centrifuged at 200rcf for 5 minutes at room temperature (RT). The supernatant was discarded, and cells were resuspended, to plate iPSCs as single cells, with 1ml of fresh medium containing mTeSR1, 50U/ml P/S, 4µg/ml Doxycycline hyclate (Sigma-Aldrich, D9891), and 10µM RI. Cells were diluted in the culture medium, and plated at a density of 25x10^3^ cells/cm^2^. Cells were cultured at 37°C with 5% CO^2^ and 20% oxygen.

#### Day 1

##### Media change

The differentiation medium was prepared containing DMEM/F12, 1%v/v N2 (Thermofisher, 17502048), 50U/ml P/S, 1%v/v MEM Non-Essential Amino Acids Solution (Thermofisher. M7145), 4µg/ml Doxycycline, 10µM RI, 10ng/ml Recombinant Human NT3 (NT3) (Peprotech, 450-03), and 10ng/ml Recombinant Human/Murine/Rat BDNF (BDNF) (Peprotech, 450-02) and filtered through a 0.22µm filter, then 0.2µg/ml Laminin was added. Cells were washed once with DPBS, and fresh prewarmed differentiation medium was added.

#### Day 2

##### Adding rat astrocytes

The second day of differentiation rat astrocytes were added to iNeurons cultures. Rat Primary Cortical Astrocytes (Thermofisher, N7745100) cultures were maintained in T75 flasks, coated with 10µg/ml Poly-D-Lysine (Sigma-Aldrich, A003-E), in growth medium composed by DMEM high glucose (Thermofisher, 11965-092), 15% Fetal Bovine Serum (FBS) (Sigma-Aldrich, F2442), 50U/ml P/S. To co-culture astrocytes with iNeurons, cells were washed once with DPBS and incubated with 5ml of TrypLE for 5 minutes at 37°C. Then astrocytes were washed with the old medium to stop the enzymatic reaction and centrifuged at 250rcf for 5 minutes. After the centrifugation, the supernatant was discarded, and the pellet was resuspended with 1ml of fresh astrocytes medium, cells were counted and plated as single cells on iNeurons at a density of 10x10^3^ cells/cm^2^. The co-culture was continued at 37°C, 5% CO^2^ and 20% oxygen.

#### Day 3

##### Media change

The differentiation medium was prepared containing Neurobasal medium (Thermofisher, 21103049), supplemented with 2% v/v B27 Supplement (B27) (Thermofiseher, 17504044), 50U/ml P/S, 1% v/v GlutaMAX™ Supplement (GlutaMAX) (Thermofisher, 35050038), 4µg/ml Doxycycline, 2µM Cytosine β-D-arabinofuranoside (AraC) (Sigma-Aldrich, C1768), 10ng/ml NT3, and 10ng/ml BDNF, the medium was then filtered through a 0.22µm filter. Cells were washed once with DPBS to remove dead astrocytes, and fresh prewarmed differentiation medium was added.

### Day 6 and 8

50% media change: From the sixth day in vitro only 50% of the medium was changed; therefore, half of old medium was discarded, and prewarmed fresh differentiation medium was added to the iNeurons culture.

The differentiation medium contained Neurobasal supplemented with 2% v/v B27, 50U/ml P/S, 1% v/v GlutaMAX, 4µg/ml Doxycycline, 10ng/ml NT3, and 10ng/ml BDNF. From day 10 on. 50% media change: old medium was discarded, and prewarmed fresh differentiation medium was added to the iNeurons culture. The differentiation medium contained Neurobasal supplemented with 2.5% FBS, 2% v/v B27, 50U/ml P/S, 1% v/v GlutaMAX, 4µg/ml Doxycycline, 10ng/ml NT3, and 10ng/ml BDNF, the medium was filtered through a 0.22µm filter.

### iNeurons lipofection

To visualize iNs soma and neurites at the single cell level, a sparse labelling method was used. Cells were plated on 13mm coverslips and lipofected at d3 with a plasmid encoding cytoplasmic GFP (pCAGG1-GFP) using the Lipofectamine™ 3000 Transfection Reagent (Thermofisher, L3000008), following the manufactures indications. Briefly, two different mixes were prepared. Mix I: 25ul OptiMEM (Thermofisher, 31985062), 1ul P3000 and 0.1ug of pCAGGs-GFP. Mix II: 25ul OptiMEM and 0.75ul Lipofectamine 3000. Mix I was added to Mix II and incubated for 15 minutes at RT. After incubation, 50µl of mix were added to iNeurons, and cells were put back in a humidified incubator set at 37°C with 5% CO2 for 48 hours. The differentiation medium was completely removed, fresh medium was added and iNs were kept in culture and fixed at different time points (d14, d21, d28, and d35) to follow neuronal maturation and synaptogenesis over time at the single cell level (see below). In parallel, non-lipofected iNeurons were kept in culture to check for neuronal markers expression.

### Cerebral organoids generation

Cerebral organoids were generated from one chimpanzee iPSCs line (SandraA) and one human iPSCs line (WTCII, obtained from Coriell institute for medical research, GM25256), following the Lancaster and Knoblich (2014) protocol(38,39). Briefly, when iPSCs reached an 80% confluency, cells were washed with PBS and detached with Accutase. iPSCs where then transferred to a falcon tube and centrifuged at 200xg for 5 minutes (Day 0). The supernatant was discarded and the cell pellet was resuspended in 1ml of fresh medium (mTeSR™1 and 10µM RI). Cells were diluted in the culture medium and plated to reach 9x10^3^ cells/well of a 96-well plate (Twin Helix, MS-9096UZ). 96-well plates were centrifuged at 280xg for 5 minutes to form Embryoid bodies (EBs) and kept in culture at 37°C, 5%CO2, 20%O2. Day 2: the medium was changed the second day in culture; old medium was discarded and fresh mTeSR™1 was added. Day 7: EBs were embedded in Geltrex and kept in culture in shaking and maintained in Differentiation medium plus B27 (DM + B27).

### Organoids electroporation

To visualize neuron architecture and spine morphology at single-cell resolution, we performed electroporation of human and chimpanzee brain organoids to achieve sparse GFP labelling.

Day 45 organoids were collected and maintained in differentiation medium lacking P/S prior to the procedure. A plasmid solution containing pCAGGS-GFP (1 µg/µL in H₂O) was prepared and kept on ice until use. Glass capillaries (borosilicate; outer diameter 1.2 mm, inner diameter 0.94 mm, 10 cm length; Sutter Instruments) were pulled using a P-1000 micropipette puller (Sutter Instruments) with the following parameters: Heat 500, Pull 60, Velocity 80, Delay 100, Pressure 280.

Organoids were transferred to PBS -/- and positioned under a dissecting microscope to identify ventricular structures. The GFP plasmid was then microinjected into individual ventricles using the prepared capillaries under visual guidance. Electroporation was performed using five pulses at 80 V, each lasting 50 ms with 500 ms intervals, delivered by an ECM830 electroporator (BTX).

Following electroporation, organoids were incubated in differentiation medium without antibiotics for 4 hours, after which they were returned to standard culture conditions. Organoids were maintained for additional 55 days in culture and then fixed (at d100) to enable subsequent reconstruction single dendrites and dendritic spines.

### MEA recordings

iNs at d3 were plated with primary astrocytes at a total cell density of 100,000 cells/well single well high-density MicroElectrode Arrays (HD-MEA) (Accura 3Brain) containing 4,096 electrodes in a 3.8 x 3.8 mm^2^ area. Spontaneous activity was recorded every 7 days, starting from d21 until d42, for 5 minutes with BioCAM DupleX (3Brain), with a sampling rate of 20,000Hz. Spike detection was performed BrainWave6 software. After spike detection spike times were used to plot Raster plots and calculate the number of active channels setting a threshold of 0.05Hz as minimum frequency to identify a channel as active.

### Organoids embedding, cryosectioning and IHC staining

Brain organoids (d100) were fixed with PFA 4% for 40 minutes and washed with PBS 1X. the organoids were then equilibrated in 30% sucrose at 4°C overnight before being embedded in TissueTek freezing medium (Leica Biosystems, Germany). The embedded samples were rapidly frozen in n-pentane (Carlo Erba). Sections of 100µm thickness were then obtained using a cryostat (Leica Biosystems) and stored at −20°C until further processing. The 100µm thick slices were then stained for 30 minutes with DAPI and then mounted with Mowiol.

### Immunofluorescence on iNs

iNeurons (lipofected and non-lipofected, based on the experimental needs) were fixed at different timepoints (d14, d21, d28, d35) with 4% paraformaldehyde (PFA) and 4% saccharose. The medium was removed, and a solution of 4% PFA and 4% saccharose was added to the dish, and cells were incubated for 8 minutes at room temperature. The fixative was discarded, and cells were washed three times with phosphate-buffered saline (PBS 1X). If necessary, fixed iNeurons were stored in PBS at 4°C for subsequent immunofluorescence analysis. After fixation, iNeurons were permeabilized with 0.05% Triton in PBS and incubated at room temperature for 10 minutes. iNeurons were quenched with 0.2N Glycine in PBS (glycine buffer) at room temperature for 30 minutes. IF buffer (0.2% Gelatin, 30mM NaCl, 0.05% Triton diluted in PBS) was used as a blocking solution. iNeurons were washed three washes times for 10 minutes at room temperature with IF buffer. Coverslips were then incubated with primary antibody diluted in IF buffer overnight at 4°C. The day after, the primary antibody solution was discarded and coverslips were washed with PBS 1X four times for 5 minutes before the incubation with the respective secondary antibody and DAPI (Thermofisher), diluted in IF buffer. The secondary antibodies were incubated for 1 hour at room temperature. After four washes with PBS (5 minutes each) the coverslips were mounted with Mowiol on a glass slide, the slides were allowed to solidify for 30minutes at room temperature and then transferred and stored at 4°C. For a comprehensive list of all antibodies used, see Table I.

### Electron Microscopy on human and chimpanzee iNs

#### Primary fixation

Cells were fixed at d28, d35, d49 and in 1,2% glutaraldehyde (Electron Microscopy Science), 66mM sodium cacodylate pH 7.4 in distilled water for 2h at room temperature, or overnight at 4°C. After fixation, cells were washed with abundant sodium cacodylate buffer 0.1M pH 7.4 at least three times for 20 minutes; to avoid the interference of glutaraldehyde with the following steps.

#### Secondary fixation or post-fixation

Cells were then post-fixed with 1% osmium tetroxide (Electron Microscopy Science), 1% potassium ferricyanide and 0.1M sodium cacodylate buffer pH 7.4 for 1h. After 2 additional washes in 0.1M sodium cacodylate buffer and 3 washes in distilled water, cells were incubated with 1% of uranyl acetate for two hours at room temperature or overnight at 4°C. Three washes with distilled water followed.

#### Dehydration

Dehydration was achieved through sequential incubation in ethanol solutions with increasing concentrations: 50%, 60%, 70%, 80%, 90%, 96% and 100%. Ethanol was diluted in distilled water and added cold (4°C), then incubated at room temperature. Cells were incubated twice with each ethanol solution for 5 minutes, except for 100% ethanol incubations that lasted for 30 minutes.

### Resin infiltration

EMbed 812 (Electron Microscopy Science) resin was used with a mixture of 1:1 for at least two hours at room temperature. Then, two additional incubations were performed, 1 hour each. For TEM processing, glass coverslips were maintained in multiwell plates untill incubation with 29Embed + Ethanol 100% 1:1. Then, they were removed and incubated with pure Embed mix placed on parafilm sheet. Resin was polymerized at 60°C for 48 hours. The portion with polymerized resin was detached from glass supports with repetitive immersion in liquid nitrogen.

#### Sectioning and TEM image acquisition

Ultra-thin sections (60-70nm) were obtained with the ultramicrotome (Leica UC 6, Leica Microsystems, Wetzlar, DE) by means of a diamond knife (Ultra 45°, Diatome) and collected on 300 mesh copper grids (Electron Microscopy Science). Sections were then stained on the grid with saturated solution of uranyl acetate in distilled water for 20 minutes and 1% lead citrate solution in distilled water. Samples were observed at the TEM (FEI Tecnai Spirit G2 and Thermo Scientific Talos L120C) at 120kV, and images were acquired with a digital camera (Olymps Megaview G2 camera and Thermo Scientific CETA 16M).

#### Image analysis

Synapses were identified by the presence of 3 synaptic vesicles in the presynaptic terminal, and a distinct synaptic cleft. The measurements of the distance of each vesicle to its closest one were obtained using the convolutional neural network-based algorithm developed by Imbrosci et al., that enables the automatic detection of synaptic vesicles in EM images (Imbrosci, Schmitz and Orlando, 2022). The synaptic surface was considered as the region comprehending the varicosity filled with 30 synaptic vesicles of the presynaptic terminal. The synaptic surface, PSD length and synaptic vesicles distance from the active zone were quantified with Fiji. Synaptic vesicles density is calculated as the ratio between synaptic vesicles number and the synaptic surface. The PSD thickness is measured as the ratio of the PSD area and the PSD length.

#### Statistical analysis

Results are reported as mean ± SEM. Statistical analyses were conducted performing either Student’s t test or one-way repeated measures ANOVA for single and multiple comparisons respectively.

### Cholera Toxin staining

iNeurons (at d28) were incubated on ice for 10 minutes, afterwards the cells were incubated on ice for 30 minutes with 1µg/ml of CholeraToxin-Alexa555 (Thermofisher, C34776) diluted in iNeurons culture medium. Cells were quickly washed twice with PBS 1X and then fixed in 4% PFA and 4% saccharose at room temperature for 8 minutes. Cholera-toxin stained iNeurons were processed by immunofluorescence with additional markers as described above.

### Synaptosomes preparation *via* differential centrifugation

For synaptosomes preparation, 1-3 million iPSCs cells were plated on a 6 multiwell plate. At d35 of differentiation cells were collected and processed to isolate the synaptosome fraction following W. B. Huttner et al., 1983 and S. Rajkumar et al., 2023 protocols (40,41). The protein concentration was then quantified in each cellular fraction with Bicinchoninic Acid assay (BCA) (Thermofisher, 23227).

### MS-based proteomics

3 million Human and Chimpanzee iNs (N=5 biological replicates) were fractionated to obtain synaptosomes fraction and processed by in-solution digestion with the PreOmics iST (PreOmics GmbH) sample preparation kit following the vendor’s instructions. 1μg of peptides were injected for each liquid chromatography-mass spectrometry (LC-MS) acquisition. The LC-MS platform consisted of a Vanquish Neo system (ThermoFisher Scientific) connected to an Orbitrap Astral mass spectrometer (ThermoFisher Scientific) operating under Tune 1.1.118.15. Mobile phases consisted of 0.1% v/v formic acid in water (mobile phase A) and 0.1% formic acid in 80% acetonitrile/water v/v (mobile phase B). The system was set up in trap-and-elute workflow with a pepmap Neo trap cartridge (thermo Scientific, 174500) The samples were separated on an EASY-Spray PepMap Neo column (75 µm x 50 cm) (ThermoFisher Scientific, ES75500PN) with a flow of 300 nl/min with a 70 minute gradient with linear separation between 5%-30% B in 60 minutes (42).

MS acquisitions were performed in narrow-window data-independent (DIA) mode (43). MS1 spectra were acquired with a resolution of 240,000 and 5ms maximum injection time with a full scan range of 380–980 m/z. Source RF lens was set to 40%.

The DIA part was performed with 300 windows of 2-Th scanning from 380 to 980 m/z under a time control of 0.6 second maximum second per duty cycle. The ion injection time was set to a maximum of 3ms per windows and the automatic gain control target to 500% (5e4 ions). The isolated ions were fragmented using HCD with 25% Normalized Collision Energy.

The resulting raw files were processed with DIA-NN 1.9.2. (43) Human and Chimp spectral libraries were generated in silico by DIA-NN from UniProt (including TrEMBL) and are included in the data deposition. Spectral libraries were generated including methionine oxidation, 1 missed cleavage, precursor charge states 1-4 and common contaminants. The 5 human and 5 chimp raw data files were analysed separately. The data was searched with a minimum peptide length of 7, 1 missed cleavage, methionine oxidation as variable modification, cysteine carbamidomethylation as fixed modification, match between runs, and the direct quantification option in DIA-NN. The data was filtered with a 0.01 q-value filter at the precursor level and 1% false discovery rate (FDR) at the protein group level.

The resulting human and chimp gene group matrices (report.gg_matrix.tsv) were merged by gene names with an inner merge. Data was filtered to minimum 80% (4/5) valid values in at least 1 group (human or chimp) and normalized using variance-stabilizing normalization (44). Missing data was imputed with the Perseus algorithm (45). Volcano plot was generated with a welch test with Benjamini-Hochberg correction for multiple testing (alpha=0.05) and a minimum fold-change filter of 1. Filtering, imputation and Volcano plotting were performed in QProMS ((4–7), https://github.com/ieoresearch/QProMS). Positive values indicate higher protein enrichment in human iNs and negative values indicate enrichment in chimpanzee neurons. For functional enrichment analysis only significantly changing proteins were considered, for a total amount of 846 proteins (447 for human synaptosomes and 399 for chimpanzee synaptosomes). GO terms (GO:BP) enrichment analysis was performed separately on the two enriched protein lists using g:Profiler (version API 1.0.0, gprofiler-official Python package). Homo sapiens annotated genome was used as statistical background. The analysis was performed with a Benjamini-Hochberg (FDR) correction for multiple testing and with an adjusted p-value < 0.05. For each species, the first higher enriched 20 terms were plotted using a Dot plot.

The correlation between proteins differentially enriched and neurological diseases was evaluated through enrichment analysis with DisGeNET library (Enrichr). The analysis was conducted on human-biased list (447 proteins) and chimp-biased proteins (399 proteins) using correction and cutoff parameters as described above. Resulting significant terms were filtered for neurological diseases terms through keyword matching.

Analysis were conducted with Python 3.12, the following packages were used pandas 2.3.2, gprofiler-official 1.0.0, gseapy 1.1.11, and matplotlib 3.10.1.

### Dot Blot

A total of 3 µg of synaptosomal proteins or 10 µL of supernatant (SN), diluted in 1× Tris-buffered saline (TBS), were directly applied onto a 0.45 µm nitrocellulose membrane (Amersham) using a vacuum filtration system (Bio-Dot apparatus, Bio-Rad).

The membrane was blocked for 1hour at room temperature (RT) with 5% (w/v) non-fat dry milk (Sigma) prepared in 1× TBS containing 0.1% Tween-20 (TBST). The membrane was then incubated with horseradish peroxidase (HRP)-conjugated cholera toxin B subunit (CHTXB) (Invitrogen), diluted in 5% milk in TBST. Incubation was performed for 1 hour at RT CHTXB was diluted 1:100,000 for synaptosomal samples, or overnight at 4 °C and diluted 1:10,000 for supernatant samples.

Following incubation, the membrane was treated with enhanced chemiluminescence (ECL; Bio-Rad) reagent for 2 min, and chemiluminescent signals were detected using a ChemiDoc MP imaging system (Bio-Rad).

Densitometric Analysis was performed using Image Lab software (version 6.1; Bio-Rad). Protein signal intensities were normalized to total protein staining assessed by Ponceau S (Sigma).

### Western Blotting

4µg of proteins from each fraction were miced with 1X NuPAGE LDS Sample buffer and NuPAGE^™^ Reducing Agent (1X final concentration, Thermofisher, NP0009). Samples were put at 70°C for 10 minutes to achieve heat-mediated denaturation. Samples were loaded on a Bolt™ Bis-Tris Plus Mini Protein 4-12% Gel (Invitrogen, NW04120BOX) in running buffer, (20X NuPAGE Tris Acetate Running Buffer LA0041 in water, LA0041) together with the precision Protein Dual Color Standards (BIORAD, #1610374) ladder. For denaturing electrophoresis, the gel ran for 1 hour at 160V constant. After the electrophoresis, proteins were transferred onto the nitrocellulose membrane using Trans-Blot Turbo Transfer System (BIORAD, 1704158) with parameters set to: 1.3A-25V for 10 minutes.

The nitrocellulose membrane was washed once with TBST for 5 min at room temperature. The membrane was then blocked with 5% milk (Merk, 70166 prepared in TTBS) for 1 hour on a shaking platform at RT. Membranes were then incubated with the primary antibody diluted in 5% milk with gentle agitation at 4°C overnight. The day after the membrane was washed for 5 min 3 times with TBST and then incubated with secondary HRP antibody diluted in TBST-5% milk for 1h at RT. Washings with TBST were then performed three times for 5 minutes each. Membranes were incubated for 5 minutes with Clarity Max Western ECL Substrate (BIORAD, 1705062) and imaged with a digital ChemiDoc imager (BIORAD).

### Bulk RNA sequencing

For RNA isolation cells were lysed using TRI-Reagent (Merck). RNA isolation was performed using Direct-zol RNA Miniprep Kit (ZYMO), according to the manufacturer’s manual. 1 ug of intact RNA was used for each RNA-seq library construction. Libraries were prepared using KAPA HyperPrep Kit (Roche); the library preparation procedure was performed according to the kit manufacturer’s instructions. For indexing, the UMI in xGen™ UDI-UMI adapters (IDT) were used. Library concentrations were measured using Quantus reader and QuantiFLUOR reagents (Promega). Library sizes were determined using TapeStation DNA ScreenTape & Reagents (Agilent) on TapeStation 4200 device (Agilent). Libraries were sequenced (2x100 bp, paired-end) using NovaSeq6000 device (Illumina).

iNs cultures were collected for RNAseq at 3 timepoints (d14, d28, and d35). RNA-seq analysis was conducted starting from gene counts obtained with featureCounts. iPSCs derived neurons and rat astrocytes were analysed separately. Human (HS) and chimpanzee (PT) iNs were processed at d14, d28, and d35; rat astrocytes samples were excluded from iNs differential analysis. For rat astrocytes comparison mono-cultures (mono) and co-cultures (co_HS or co_PT) were compared at all timepoints. For further analysis the Ensembl version suffix was deleted.

Counts quality was evaluated by library dimension and log2(CP; + 1) values distribution both calculated with edgeR::cpm(…, log = TRUE, prior.count = 1). Mitochondrial and ribosomal genes were removed based on gene symbol. Scarcely expressed genes were also excluded, genes were kept only when showing at least 10 raw counts in at least two samples.

### TPM data normalization

TPM (Transcripts Per Million) values were calculated by dividing count per gene length (in kilobases from featureCounts), then each sample was normalized on a million transcripts. This calculation was performed on the complete matrix before the removal of mitochondrial and ribosomal genes. TPMs were only used for descriptive analysis, namely cellular markers check, synaptic makeup, cholesterol pathway, and GM1 metabolism related genes panels. TPMs were not used for differential expression analysis.

### Differential analysis by DESeq2

Differential expression analysis was performed on filtered (with DESeq2) raw counts. This workflow models count data with a negative binomial distribution, it estimates sequencing depth normalization factors and applies a moderated estimation of dispersion. The model coefficients were evaluated through Wald test with DESeq2 predefined settings. For iNeurons the following model was used:

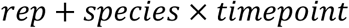

For rat astrocytes the following model was used:

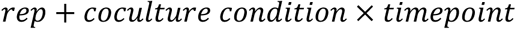

Every replicate was taken into consideration to correct the systematic effect associated to each plate. The comparison between HS and PT neuronal enriched genes was perfomed at d14, d28, and d35; the same timepoints were used for differential gene expression analysis in rat astrocytes. Astrocytes functional enrichment was performed only at d35. p-values were corrected with Benjamini-Hochberg (FDR) correction. Then, for a better comparison, log2 fold change was shrunk by ashr function. Gene were classified as differentially expressed if the subsequent thresholds were satisfied:

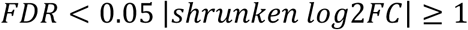

### Principle Component Analysis

For Principle component Analysis (PCA) and sample distances heatmap variance stabilizing transformation was applied:

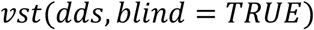

PCA was calculated on 1 thousand genes with the highest variance after VST transformation. Sample distances were calculated on VST matrix through Manhattan distance:

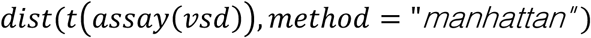

Sample distances heatmap was organized coherently to metadata, and annotated for species, condition, and timepoint.

### Gene Ontology analysis

Gene Ontology (GO) analysis was performed on differentially expressed genes at d35 for human, chimpanzee iNs, mono-cultures and co-cultures of rat astrocytes.

For iNeurons two lists were generated human enriched genes and chimpanzee enriched genes. Functional annotations were done using gprofiler2::gost(), GO:BP, with FDR correction and FDR treshold < 0.05.

For iNs GO analysis, g:Proflier was used and for both human and chimpanzee enriched genes, and organism = “hsapiens” was used. For astrocytes functional analysis organism = “rnorvegicus” was used. The specified organism defines the GO annotation database, instead the statistical background was separately defined. As a background was used a personalized one equivalent to tested and annotated genes from the differential analysis. This was specified in g:Profiler through custom_bg = universe e domain_scope = “custom”, so that non detected genes related biases were reduced.

For astrocytes (at d35) three lists were analysed: enriched genes (i) in co_HS (ii) in co_PT compared to mono-cultures, and (iii) in mono-cultures.

Lipid associated biological processes were obtained filtering significant GO:BP terms produced by g:Profiler by using predefined key words related to “lipids”, “sterols”, “lipoproteins”, “phospholipids”, “sphingolipid”, “glicosphingolipids”, “fatty acids”, and “lipid droplets”.

Enrichment analysis in gene-pathology association was evaluated on differentially expressed neuronal genes at d35 using DigGeNET through enrichR. Only terms showing Adjusted.P.value < 0.05 were kept, and subsequently filtered for neurological pathologies

### Proteomics comparison

Fora each neuronal sample eight synaptic signatures were evaluated: pre-synaptic compartment, excitatory post-synaptic compartment, inhibitory post-synaptic compartment, glutamatergic identity, GABAergic identity, synaptic adhesion, synaptic vesicles cycle, and maturation markers. Each signature score is the mean of log2(TPM + 1) values of included genes. For each signature and timepoint (d14, d28, d35) HS and PT comparison were performed with one-way ANOVA:aov(score ∼ species). The resulting p-values were corrected with Benjamini-Hochberg method. Cholesterol and GM1 panels were plotted as heatmaps of DEGs in HS and PT. Positive values indicated a higher expression in PT, and negative values indicate a higher expression in HS.

RNA-seq and proteomics concordance was evaluated on d35 neurons on shared gene simbols. When more than one Ensembl annotation were associated to the same gene simble, only the one with higher baseMean was selected. The log2 fold change HS vs. PT global concordance was evaluated with Spearman correlation.

### Software

RNA-bulk seq analysis were performed with R 4.6.0., using the following packages DESeq2, edgeR, ashr, AnnotationDbi, org.Hs.eg.db, org.Rn.eg.db, gprofiler2, enrichR, ggplot2, pheatmap, openxlsx.

---

**Supplementary Figure 1.**
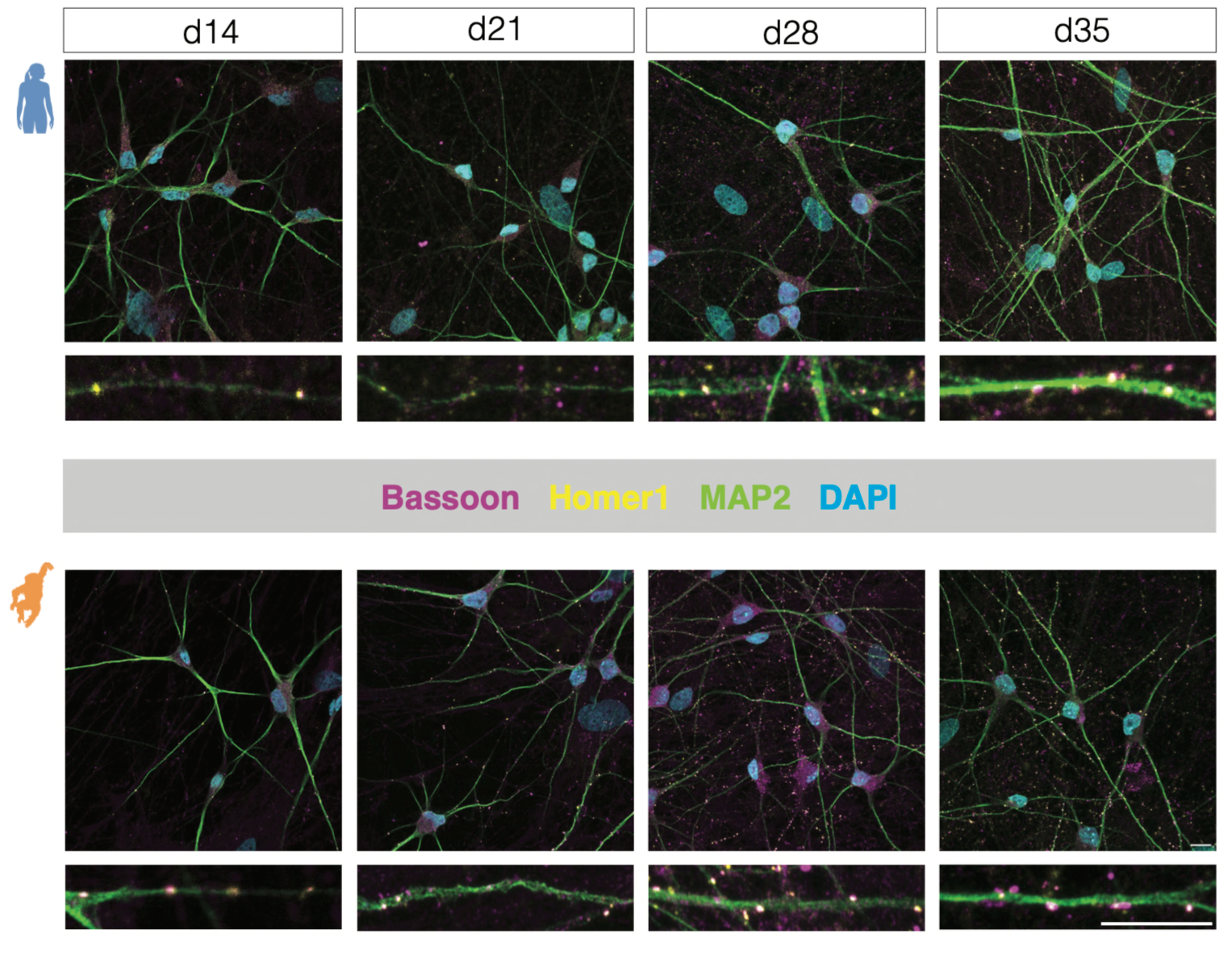
Time course of synapse formation in human and chimpanzee induced neurons. Representative immunofluorescence images of human (top) and chimpanzee (bottom) induced neurons (iNs) at d14, d21, d28, and d35 of differentiation. Neurons were stained for the presynaptic marker Bassoon (magenta), the postsynaptic marker Homer1 (yellow), the neuronal marker MAP2 (green), and DAPI (cyan). Higher-magnification views of dendritic segments are shown below each image to illustrate the progressive accumulation and colocalization of pre- and postsynaptic puncta during maturation. Scale bars 10µm. Both human and chimpanzee neurons progressively acquire Bassoon-positive and Homer1-positive puncta over time, indicating ongoing synapse formation throughout the differentiation process. Representative images correspond to the quantifications shown in Figure 3.

**Supplementary Figure 2.**
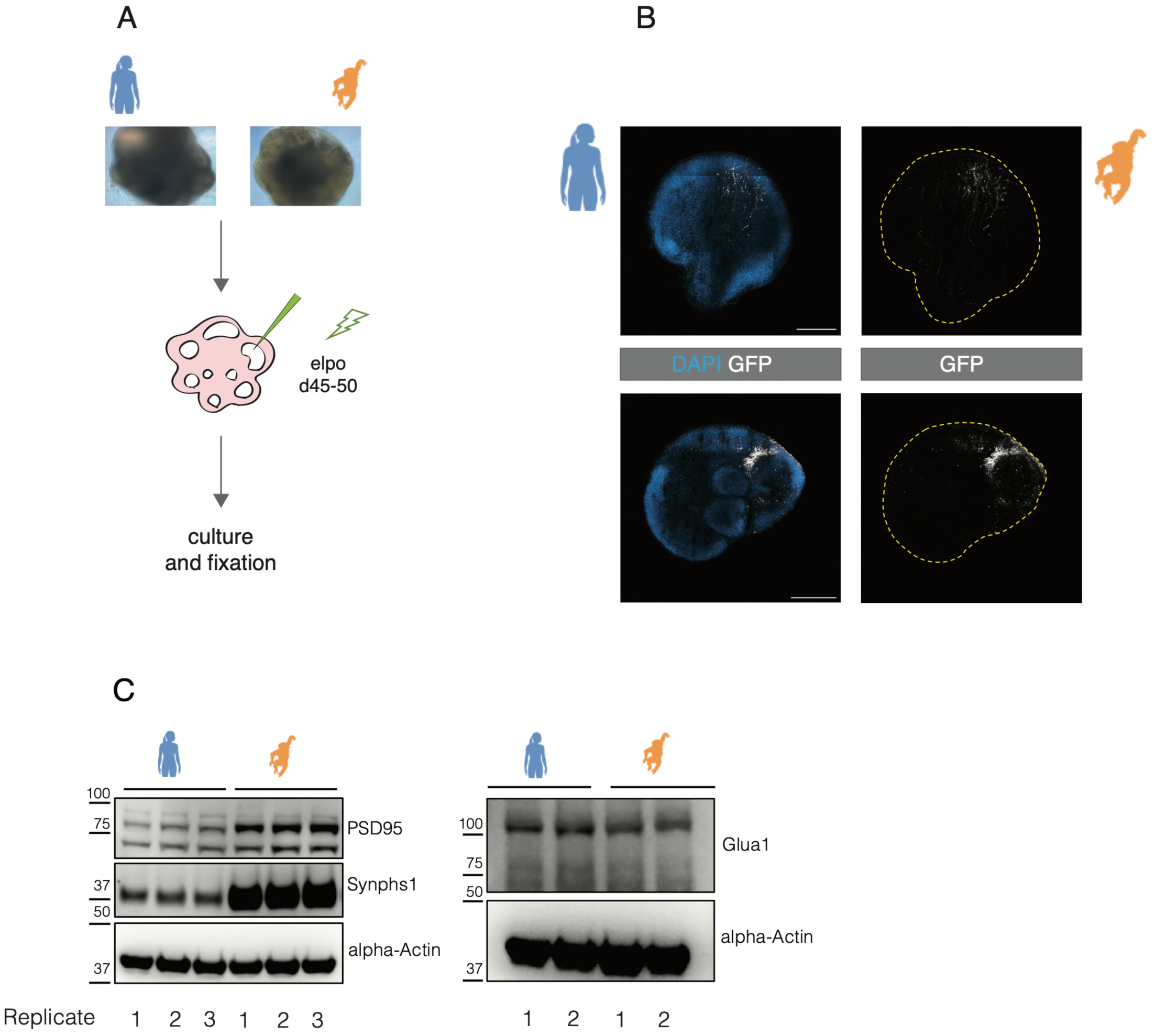
Characterization of chimpanzee and human cerebral organoids and validation of synaptic marker expression. (A) Experimental workflow for sparse neuronal labeling in cerebral organoids. Human and chimpanzee organoids were electroporated with a GFP-expressing plasmid between days 45 and 50 of differentiation and maintained in culture prior to fixation and analysis. (B) Representative images of electroporated human (top) and chimpanzee (bottom) cerebral organoids. GFP-positive cells identify sparsely labeled neurons within organoid tissue. DAPI labels nuclei. Dashed outlines indicate organoid boundaries. Scale bars, 1mm. (C) Immunoblot analysis of synaptic protein expression in human and chimpanzee induced neuronal cultures. PSD95, Synaptophysin1, and GluA1 are detected in independent biological replicates from both species. α-Actin was used as loading control.

**Supplementary Figure 3.**
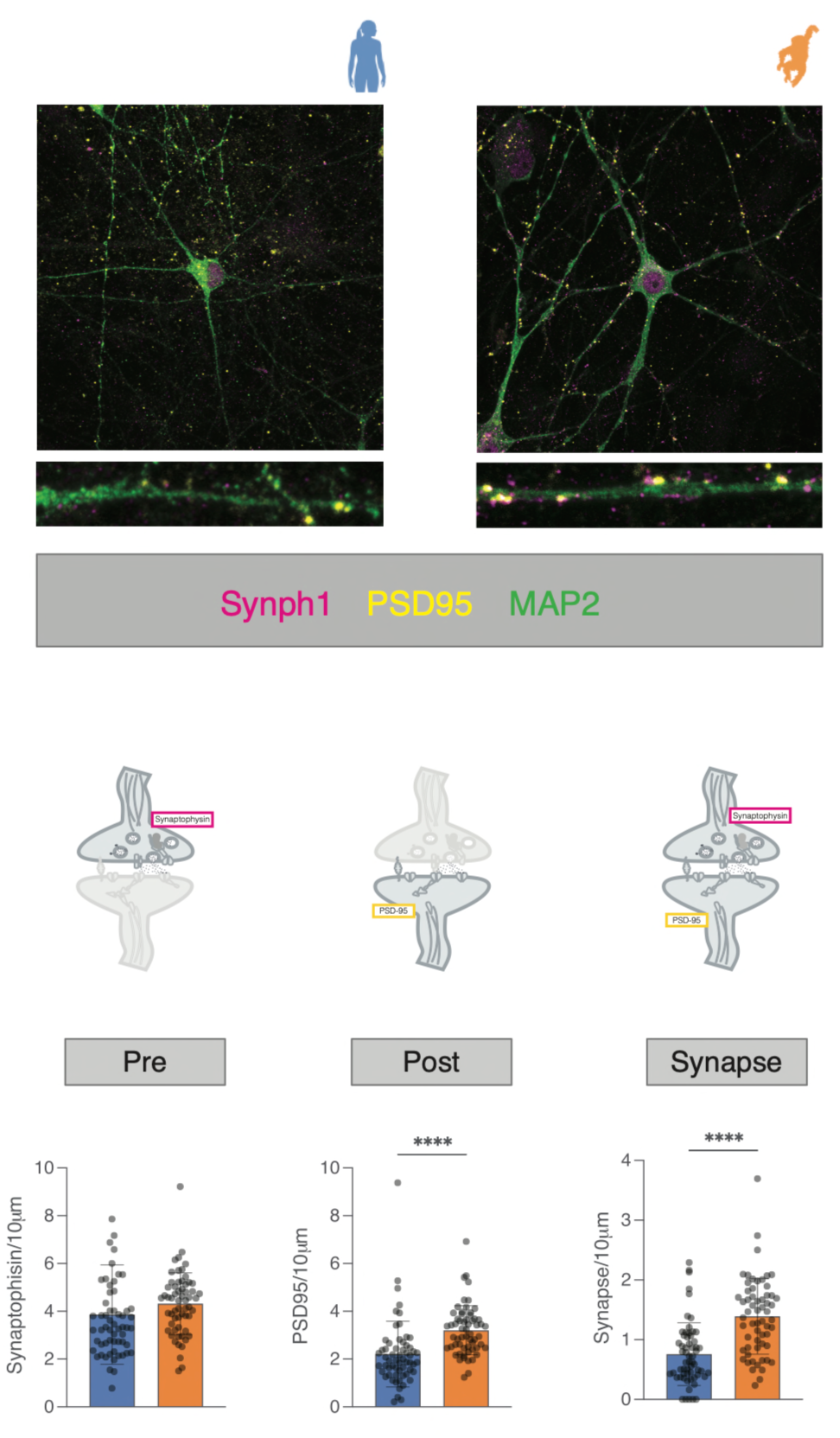
Independent validation of synapse quantification using Synaptophysin1 and PSD95. Representative immunofluorescence images of human (left) and chimpanzee (right) induced neurons stained for the presynaptic marker Synaptophysin1 (magenta), the postsynaptic marker PSD95 (yellow), and the neuronal marker MAP2 (green). Higher-magnification views of dendritic segments are shown below each image. Schematic representation of the quantification strategy used to identify presynaptic puncta (Synaptophysin1-positive), postsynaptic puncta (PSD95-positive), and putative synapses defined as juxtaposed pre- and postsynaptic puncta. Quantification of Synaptophysin1-positive puncta, PSD95-positive puncta, and colocalized synaptic puncta normalized to dendritic length. Consistent with Bassoon/Homer1 measurements shown in Figure 3, chimpanzee neurons exhibit increased densities of postsynaptic puncta and synaptic contacts relative to human neurons, whereas presynaptic puncta density is comparable between species. Each dot represents an individual dendritic segment. A total of 50-60 dendrites were analyzed for each replicate, 1 technical replicate were performed, 1 biological replicate. Statistical significance was determined using Welch’s t test, bars indicate mean ± [SD]. ****P<0.0001.

**Supplementary Figure 4.**
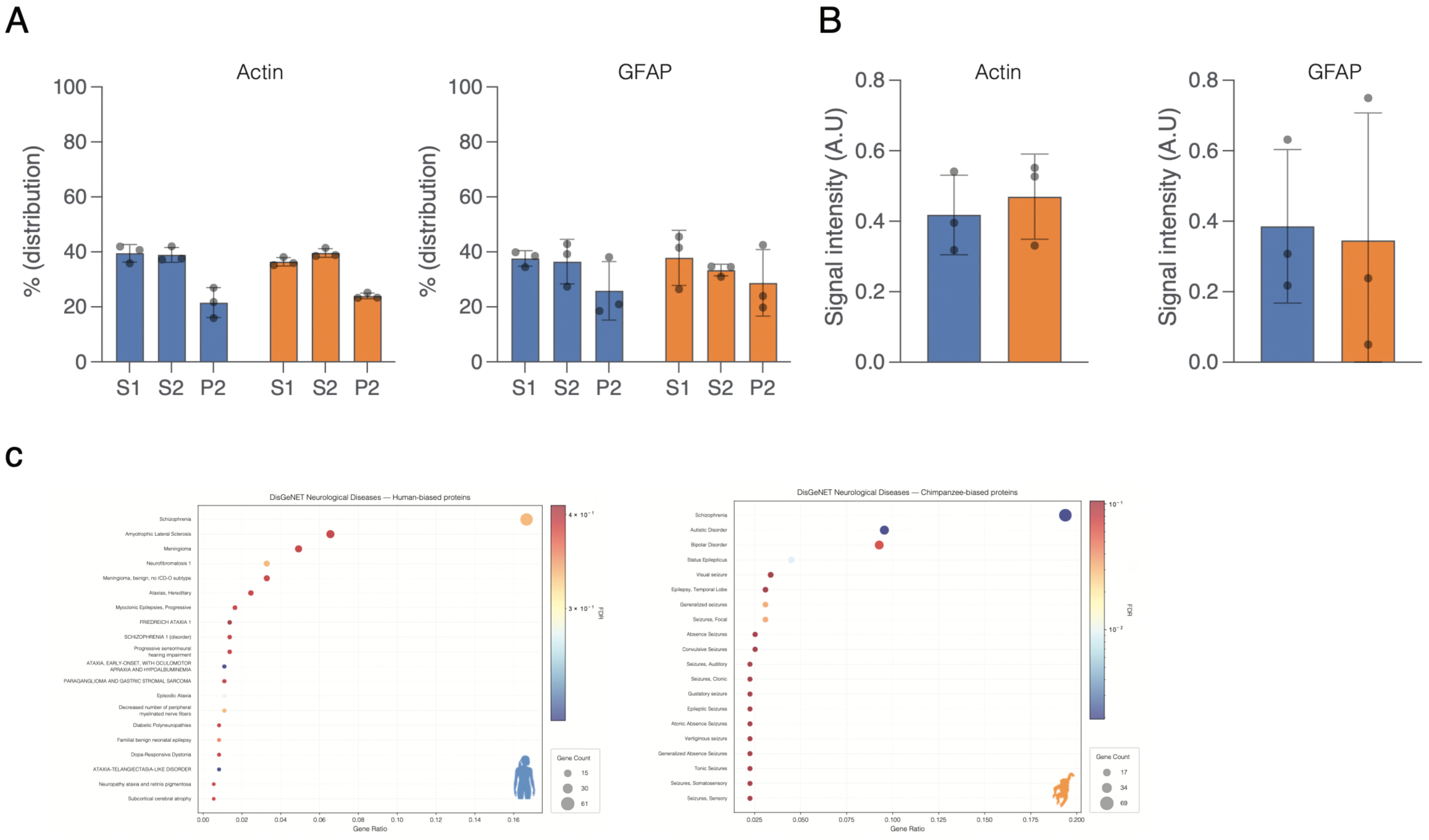
Additional proteomics characterization of human and chimpanzee neurons. (A) Quantification of protein enrichment across fractions (S1, S2, P2) for Actin, GFAP. A total of 3 technical replicates were performed, 1 biological replicate. Statistical significance was determined using two-way ANOVA (multiple comparisons), bars indicate mean ± [SD]. (B) Quantification of protein levels (actin, GFAP) in the P2 fraction. A total of 3 technical replicates were performed, 1 biological replicate. Statistical significance was determined using Welch’s t test, bars indicate mean ± [SD]. (C) Gene–disease enrichment analysis (DisGeNET) reveals that human-enriched proteins overlap with gene sets linked to neurological and psychiatric disorders, whereas chimpanzee-enriched proteins are preferentially associated with synaptic function-related pathways. These associations reflect overlap with known gene networks and do not imply causal relationships. A total of 5 technical replicates were performed, 1 biological replicate.

**Supplementary Figure 5.**
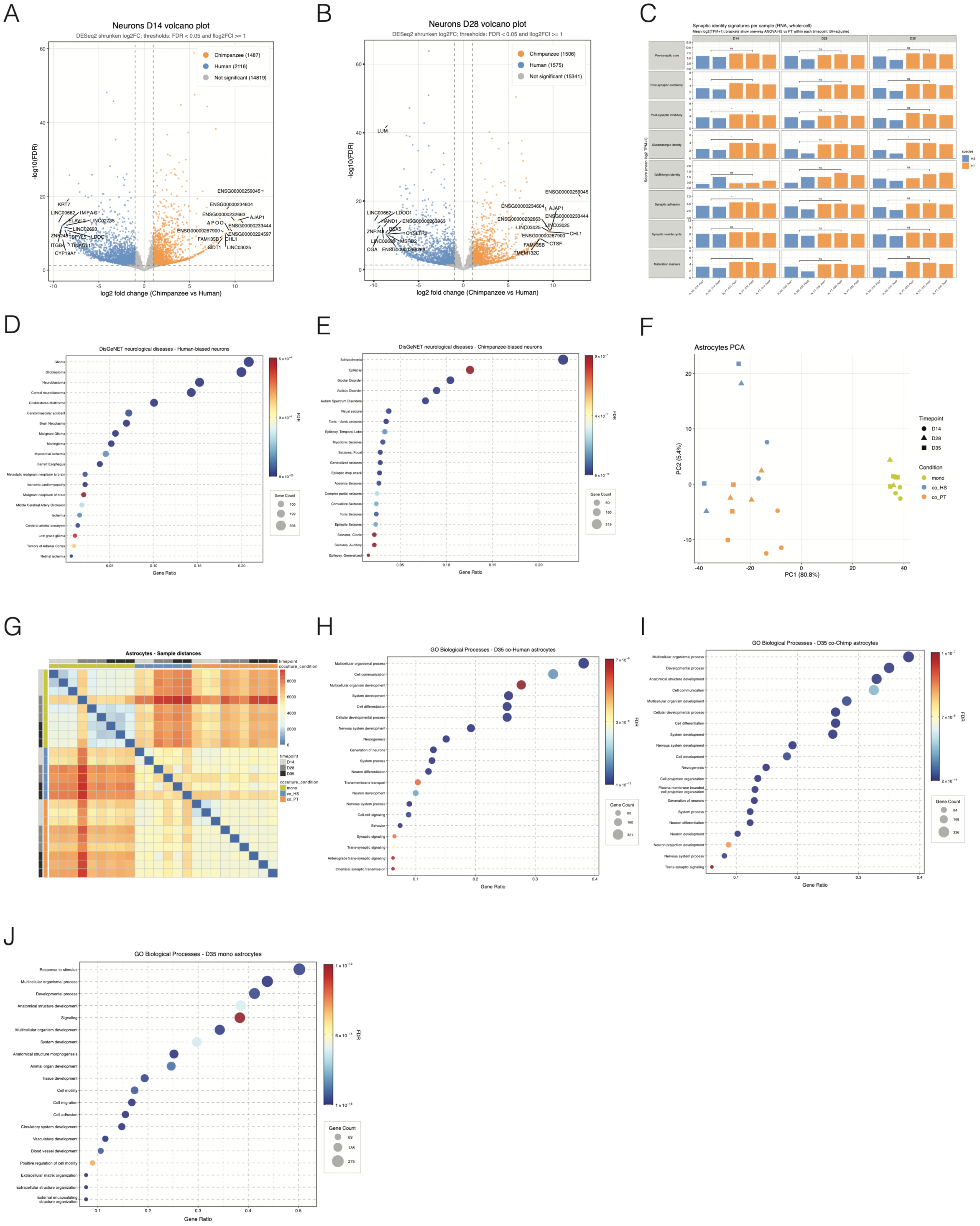
Additional transcriptomic characterization of human and chimpanzee neurons and astrocytes. (A-B) Volcano plots showing differential gene expression between human and chimpanzee neurons at d14 (A) and d28 (B). Genes enriched in human neurons (blue) and chimpanzee neurons (orange) are shown according to fold change and statistical significance. (C) Enrichment analysis of genes associated with synaptic density signatures across developmental stages and species. Relative enrichment scores are shown for selected synaptic and neuronal gene sets. Statistical significance was determined using one-way ANOVA. (D-E) DisGeNET enrichment analysis of neurological disease-associated gene sets represented among genes preferentially expressed in human (D) and chimpanzee (E) neurons. Dot size indicates gene count and color indicates adjusted significance values. (F) Principal component analysis (PCA) of transcriptomic profiles from rat astrocytes cultured alone or co-cultured with human or chimpanzee induced neurons. Samples segregate according to culture condition, indicating substantial transcriptional remodeling of astrocytes in response to neuronal co-culture. (G) Sample-to-sample distance heatmap of astrocyte transcriptomes. Replicates cluster according to culture condition, with clear separation between astrocyte monocultures and astrocytes maintained in neuronal co-culture. (H–J) Gene Ontology (GO) biological process enrichment analysis of astrocyte transcriptomes. Enriched biological processes identified in astrocytes cultured alone (H) and in astrocytes co-cultured with human (I) or chimpanzee (J) neurons are shown. Co-cultured astrocytes exhibit enrichment for pathways associated with cell–cell communication, transmembrane transport, signaling, and synaptic-related processes. Bubble size indicates gene count and color indicates adjusted significance values (FDR). Differential expression analyses were performed using DESeq2 with Benjamini–Hochberg correction for multiple testing.

